# S100A6 gliotransmission modulates neuronal proteostasis and energy production in the mouse and human brain

**DOI:** 10.64898/2026.04.10.717619

**Authors:** Valentina Cinquina, Erik Keimpema, Alán Alpár, Alexei Verkhratsky, Tibor Harkany

## Abstract

Calcyclin binding protein (CaCyBp) is an evolutionarily conserved protein that converts S100 protein family-derived signals to protein turnover through an ubiquitin ligase interaction domain within its C-terminus. Despite its predicted roles in cellular differentiation, CaCyBp expression alone and in relation to S100A6, its prototypic ligand, during fetal and postnatal organ development remains incompletely understood. Here, we combined cell-resolved neuroanatomy and biochemistry to reveal not only organ system-level CaCyBp expression but also its region-, circuit-, and neuron-specific distribution in the mouse brain from embryonic day 18.5. We extended these studies to human brains past gestational week 27, corroborating that neurons, but not astrocytes, microglia or oligodendrocytes, harbored CaCyBp. Conversely, S100A6 was detected only in astrocytes and ependymoglia in both species. Ultrastructural studies localized CaCyBp to mitochondria in neurons, with genetic loss-of-function and recombinant S100A6-induced signaling implicating CaCyBp in the regulation of the mitochondrial electrochemical gradient controlling ATP production. Overall, pleiotropic CaCyBp expression in neurons suggests a maintenance receptor-like role impacting cellular respiration, thus affecting proteostasis upon glia-derived S100A6 signals.

## Introduction

Calcyclin binding protein, also known as ‘seven in absentia homolog 1’ (Siah-1)-interacting protein (CaCyBp/SIP), is an evolutionarily conserved multidomain protein that regulates cell proliferation^1^, differentiation, and survival^2,3^ in many tissues^4^ through its structurally-distinct N-terminus, central (CS) and non-structured C-terminal SGS regions^5^. These domains allow for differential binding to protein targets, including Siah-1, S-phase kinase-associated protein 1 (Skp1) in the C-terminus^6,7^, tubulin in the N-terminus^8^, and heat shock protein 90^9–11^ at an as yet unknown site. More so, CaCyBp also exhibits intrinsic phosphatase activity with its targets including extracellular signal-regulated kinases (ERK1/2), and p38 mitogen-activated protein kinase (MAPK)^12^. CaCyBp activity is likely ligand dependent because its binding to calcyclin (S100A6) reduces downstream β-catenin degradation^13^, ERK1/2 dephosphorylation^12^ along with diminished phosphatase activity towards cytoskeletal proteins^14^.

Both Siah-1 and Skp1 are notable drivers of tumorigenesis^15,16^. It is therefore that CaCyBp has been studied in the context of cancer pathobiology with its dysregulated levels considered as prognostic markers for cancer progression^17^, invasiveness^18^, and metastasis^19^, in, e.g., neuroblastomas^20,21^. Nevertheless, this impactful knowledge on CaCyBp in disease states^17,22–24^ is insufficiently mirrored by studies on CaCyBp physiology, particularly organ development, and cellular differentiation within, including the brain.

Recently, we have shown that S100A6 expressed in and secreted from astrocytes mediates a novel form of gliotransmission during brain development^10^. Once released, S100A6 interacted with CaCyBp in neurons to modulate their morphogenesis and synaptic connectivity^10^. The molecular mechanism how S100A6 signaling could slow neuritogenesis by impacting cellular bioenergetics remained unexplored. We have closed this gap by using quantitative cell-resolved neuroanatomy to localize CaCyBp along with its ligand, S100A6, in both the mouse and human central nervous systems. We found ubiquitous and early CaCyBp expression in neurochemically heterogeneous classes of postmitotic neurons. In contrast, S100A6 was exclusively localized to astrocytes in brain regions rich in CaCyBp. At the ultrastructural level, a portion of CaCyBp localized to mitochondrial membranes where its stimulation affected the maintenance of the electrochemical gradient generated by proton pumps during oxidative phosphorylation. Thus, we suggest that S100A6-CaCyBp signaling between glial cells and neurons is relevant to neuronal maturation and life-long survival^10^.

## Results

### Body-wide CaCyBp distribution in human and mouse

The cell-resolved distribution of *Cacybp* mRNA and CaCyBp protein during fetal organ development is unknown, except in the cerebral cortex of the rodent brain^23^. Therefore, we first mapped *Cacybp* mRNA and protein in organ systems of both mouse and human. As a point of reference, we collated *Cacybp* gene expression data from the Human Protein Atlas (https://www.proteinatlas.org/ENSG00000116161-CACYBP)^25^ of tissues in the adult human body. This captured *Cacybp* mRNAs in all organs studied, with a ‘top five’ distribution at brain > eye > reproductive tract > muscle > bone marrow (Supplementary Fig. 1A,2A).

Next, body-wide anatomical screens were performed in fetal mice at embryonic day (E)18.5 with an anti-CaCyBp antibody that has been extensively validated (Supplementary Fig. 2B-C)^10,25^. CaCyBp immunoreactivity was found in the brain, spinal cord, dorsal root ganglia, digestive tract, and thymus (Fig. 1A-H; Supplementary Fig. 1C), reminiscing human expression data. Within the fetal brain, somatodendritic CaCyBp immunoreactivity invariably characterized all areas. qPCR and Western blotting corroborated broad CaCyBp distribution in all areas analyzed at both E18.5 and postnatal day (P)120 (Fig. 1I-I_2_; Supplementary Fig. 2D). Thus, one can propose a ‘maintenance factor’-like role for CaCyBp, rather than being specialized for signaling in temporally-refined contexts^10^.

**Figure 1.**
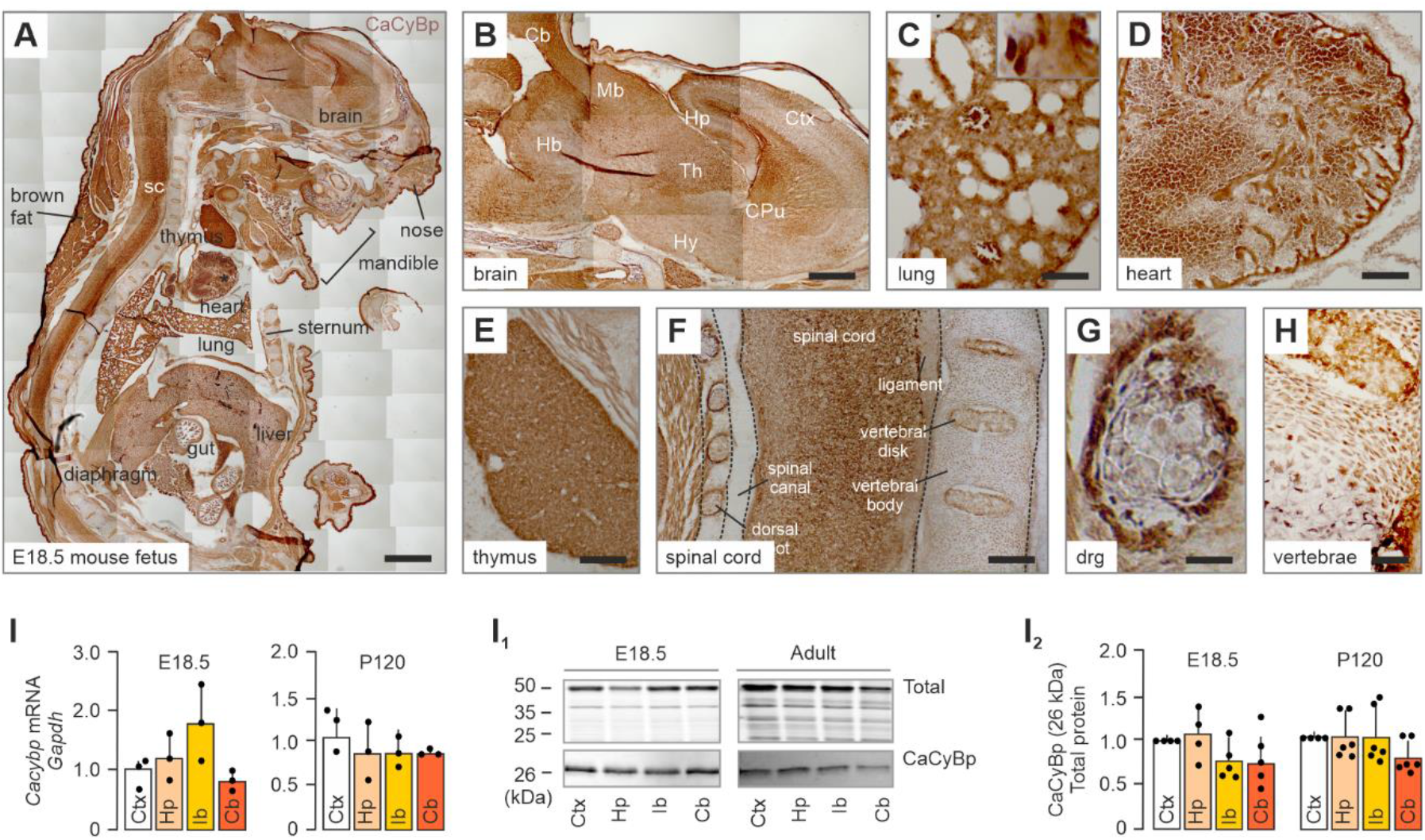
Body-wide CaCyBp localization in the developing mouse. (**A-H**) Chromogenic amplification of CaCyBp immunoreactivity (brown) in a whole-body section of an E18.5 mouse fetus, showing expression in brain (B), lungs (C), heart (D), thymus (E), spinal cord (F), dorsal root ganglia (G), and within the vertebrae (H). (**I-I**_**2**_) Bulk quantitative mRNA (N) and protein analyses (I,I_1_) of CaCyBp showed no significant differences between brain regions at E18.5 and adulthood (P120). Bar graphs represent means ± s.e.m. *Abbreviations*: Cb, cerebellum; Ctx, cortex; Hp, hippocampus; Hy, hypothalamus; Ib, interbrain (hypothalamus + thalamus); Mb, midbrain; sc, spinal cord; Th, thalamus. For antibody validation see **Supporting Fig. 1**. *Scale bars* = 1 mm (A); 500 µm (B,F); 100 µm (C-G).

### CaCyBp in fetal mouse and human brains

Fluorescent *in situ* hybridization (FISH) and immunohistochemistry mapped CaCyBp mRNA and protein, respectively, at cellular resolution in E18.5 mouse brain. Moderate-to-high signal intensity was seen in cell-dense areas, with white matter being devoid of labeling (Supplementary Fig. 3A). In the dorsal forebrain, CaCyBp immunoreactivity particularly accumulated in proliferative radial glia of the lateral ventricle (Fig. 2A,A_1_), with progeny radiating towards both the cerebral cortex and striatum (Fig. 2A_2_). To verify that CaCyBp-like immunoreactivity dominated in postmitotic neuroblasts during their radial migration, 5-ethynyl-2′-deoxyuridine (EdU, a thymidine analog)^26^ was administered in pregnant dams intraperitoneally at the peak of cortical neurogenesis^27^ (E14.5). At E18.5, cells triple-labeled for EdU, NeuN (prototypic neuronal marker)^28^, and CaCyBp were found in the subventricular zone and deep cortical plate (Fig. 2A_1_). This notion was supported by FISH indiscriminately co-localizing *Rbfox3*, encoding NeuN^29^, and *CaCyBp* mRNAs across the cortical plate (Fig. 2B). Similarly, CaCyBp was broadly expressed in all subfields of the hippocampal formation (Fig. 2C). CaCyBp immunoreactivity dominated in cell-dense layers, e.g., the prospective Cornu Ammonis (CA)1-3 subfields and granule cell layer of the dentate gyrus (Fig. 2C_1_-C_3_). Dual FISH for *Rbfox3* and *CaCyBp* mRNAs confirmed neuronal expression (Fig. 2D). In the hypothalamus, CaCyBp labelling was particularly significant in the suprachiasmatic and arcuate nuclei, with CaCyBp immunoreactivity interspersed, but not co-localized, with vimentin, a cytoskeletal filament in dorsal ependyma and tanycytes^30^, both of glial origin (Supplementary Fig. 3B,C). In the midbrain, CaCyBp immunoreactivity decorated the substantia nigra *pars compacta* but was absent from the substantia nigra *pars reticulata* (Supplementary Fig. 3D).

**Figure 2.**
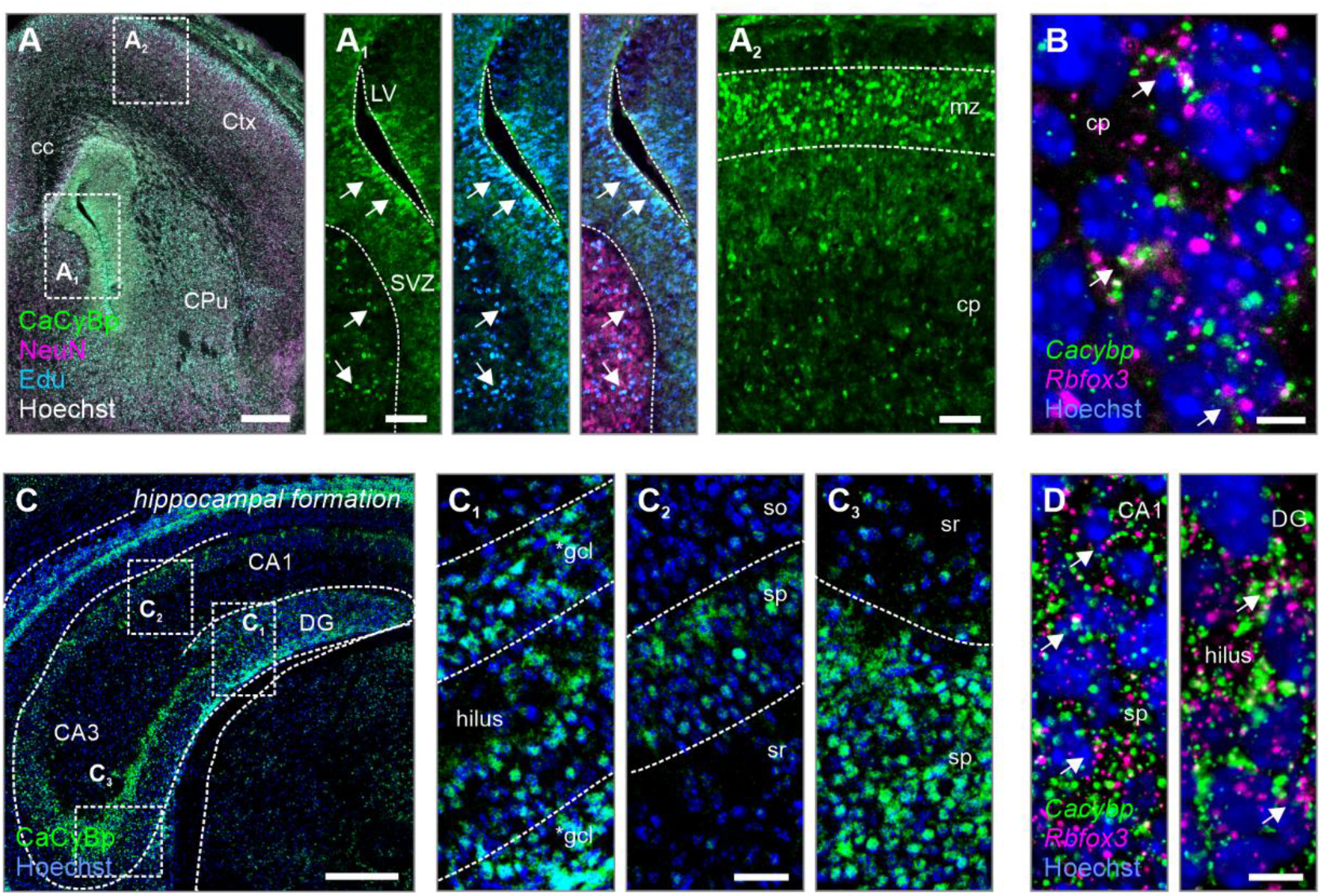
CaCyBp in the developing mouse forebrain. (**A-A**_**2**_) CaCyBp protein expression in the ventricular zone (A_1_; *arrows denote co-localization*) and cortical layers (A_2_). (**B**) *In situ* hybridization with *Rbfox3* and *Cacybp* in the neocortex. (**C-C**_**3**_) CaCyBp expression along the CA1-CA3 subfields and dentate gyrus of the hippocampus. (**D**) Multi-colour *in situ* hybridization with *Rbfox3* and *Cacybp* in the dentate gyrus. *Abbreviations*: 3V, third ventricle; Aq, cerebral aqueduct; CA1-3, cornu Ammonis; cc, corpus callosum; cp, cortical plate; CPu, caudate putamen; Ctx, cortex; DG, dentate gyrus; f, fimbria; *gcl, immature granular cell layer; Hp, hippocampus; Hy, hypothalamus; lot, lateral olfactory tract; LV, lateral ventricle; mz, marginal zone; SN, substantia nigra; sp, stratum pyramidale; spt, septum; sr, stratum radiatum; SVZ, subventricular zone; Th, thalamus. *Scale bars* = 200 µm (A), 200 µm (C), 50 µm (A_1_,A_2_,C_1_,C_2_,C_3_), 20 µm (B,D).

CaCyBp immunoreactivity in gestational week 27-34 human fetuses (early third trimester; Fig. 3A)^31,32^ also labeled cortical and subcortical areas (Fig. 3B-B_3_), as well as proliferative zones, with the medial ganglionic eminence (MGE) exhibiting the highest labeling density (Fig. 3C-C_2_). In the hippocampal formation, CaCyBp immunoreactivity accentuated in cell-and dendrite-dense layers of all CA subfields (Fig. 3D-D_3_). In subcortical areas, large-diameter somata with CaCyBp immunoreactivity were scattered in the putamen, globus pallidus, and neighboring claustrum (Fig. 3E; Supplementary Fig. 4A,B), as much as in the thalamus, hypothalamus, and substantia nigra *pars compacta* (Supplementary Fig. 4C-E). These data highlight conserved regional, and (sub-)cellular distribution for CaCyBp in fetal mouse and human brains.

**Figure 3.**
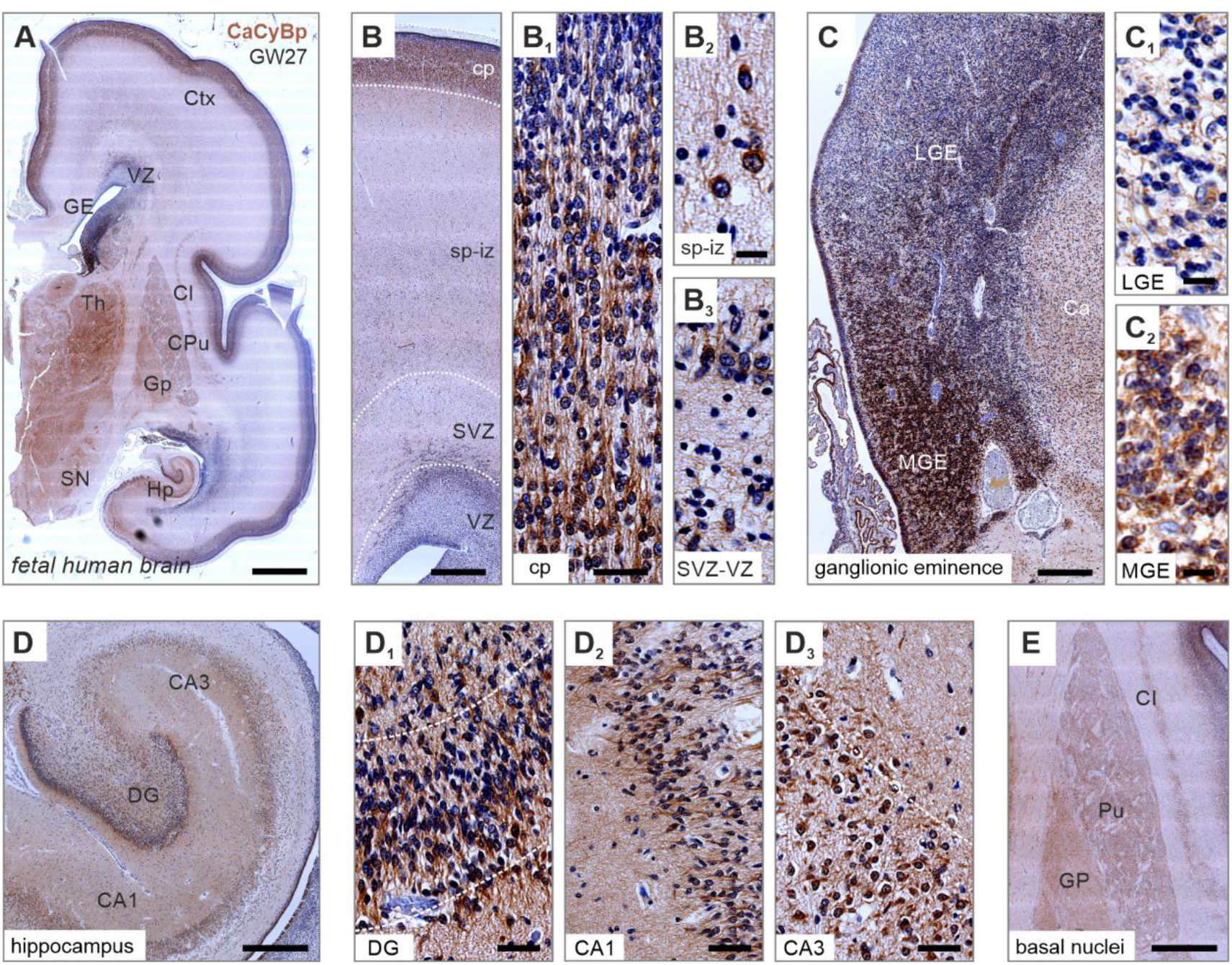
Expression of CaCyBp in the fetal human brain. (**A**) Histochemical overview of CaCyBp expression in the human brain at gestational week 27 (GW27). (**B-C**_**2**_) CaCyBp protein expression in the cortex with high levels in the cortical plate (B_1_), and ganglionic eminence (C-C_2_). (**D-D**_**3**_) In the hippocampal formation, CaCyBp was detected in the granular layer of the dentate gyrus (D_1_) and the *stratum pyramidale* of the CA subregions (D_2_,D_3_). (**E**) CaCyBp expression in the globus pallidus and putamen. *Abbreviations*: CA1-3, cornu Ammonis 1-3 subfields; Cl, claustral complex; cp, cortical plate; CPu, caudate putamen; Ctx, cortex; DG, dentate gyrus; GE, ganglionic eminence; Hp, hippocampus; Hy, hypothalamus; LGE, lateral ganglionic eminence; MGE, medial ganglionic eminence; PU, putamen; SN, substantia nigra; sp-iz, subplate intermediate zone; SVZ, subventricular zone; Th, thalamus; VZ, ventricular zone. *Scale bars* = 1 mm (A), 200 µm (C,D,E), 40 µm (B), 20 µm (B_1_,D_1_,D_2_,D_3_), 10 µm (B_2_,B_3_,C_1_,C_2_).

### CaCyBp immunoreactivity in adult mouse and human brains

The localization of CaCyBp immunoreactivity in adult tissues from both mice (P120; Fig. 4A-C_3_) and humans (57-63 years; Fig. 4D-E_3_) were addressed next. In the cerebral cortex, CaCyBp somatic labeling exhibited a dorsal-to-ventral increase in density (∼60% in layers V/VI; Fig. 4A-B_1_). In the hippocampus, CaCyBp immunoreactivity labelled the *strata pyramidale* and *radiatum* of the CA subfields, dendrites in the CA3 *stratum lucidum*, with some perikarya also scattered in the *strata oriens* and *laconosum moleculare* (Fig. 4C-C_2_; Supplementary Fig. S5A-B_1_). In the DG, CaCyBp immunoreactivity was seen in the subgranular zone and polymorphic layer/hilus (Supplementary Fig. S5B_2_). Within the subgranular zone, CaCyBp immunoreactivity was prominent in neuroblasts (type 2/3 cells; Supplementary Fig. S6A) positive for doublecortin, a developmentally-expressed microtubule-associated protein^33^, as much as in immature neurons immunoreactive for calretinin, an EF-hand Ca^2+^-binding protein^34^. The molecular layer only contained occasional cells. The granule cell layer was rich in terminal labeling, likely from recurrent axons of hilar mossy cells (Fig. 4C_3_). CaCyBp immunoreactivity was absent from glial fibrillary acidic protein (GFAP)-positive astrocytes (Supplementary Fig. S6B)^35^, supporting that CaCyBp expression is limited to neuronal lineages (positive for doublecortin, calretinin and calbindin D28k; Supplementary Fig. S6B_1_-B_3_). This CaCyBp distribution is a postnatal feature which existed by P6 (Supplementary Fig. S6C-C_2_) and likely indicates activity-dependent enrichment.

**Figure 4.**
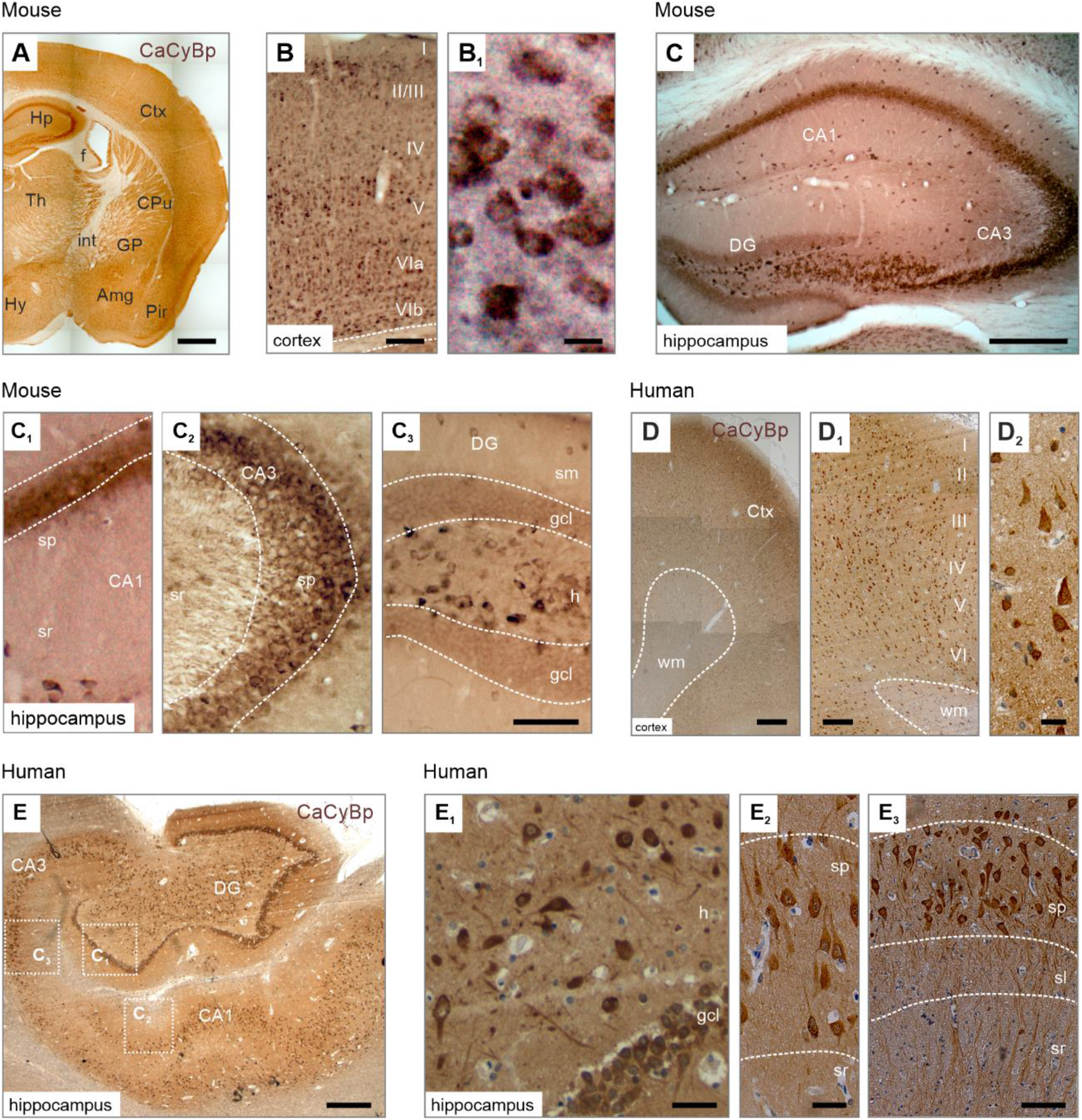
Expression of CaCyBp in the adult mouse and human forebrain. (**A-B**_**1**_) CaCyBp protein in the cerebral cortex accumulated in deep cortical layers (B). (**C-C**_**3**_) In the mouse hippocampal formation, CaCyBp was detected in the *stratum pyramidale* of the CA1-3 subfields (C_1_,C_2_), as well as the hilus of the dentate gyrus (C_3_). (**D-D**_**2**_) CaCyBp localized to cells across all cortical layers in the human cortex. (**E-E**_**3**_) In the human hippocampal formation, CaCyBp was found expressed in the granular and polymorphic layers of the dentate gyrus (C_1_), as well as in the *stratum pyramidale* of the CA1-CA3 subfields (C_2_, C_3_). *Abbreviations*: Amg, amygdala; CA1-3, cornu Ammonis; CPu, caudate putamen; Ctx, cortex; DG, dentate gyrus; gcl, granular cell layer; GP, globus pallidus; h, hilus; Hp, hippocampus; Hy, hypothalamus; sl, stratum lacunosum moleculare; sp, stratum pyramidale; sr, stratum radiatum; Th, thalamus; wm, white matter.. *Scale bars* = 500 µm (A), 200 µm (D,E), 100 µm (C), 50 µm (B,C_1_,C_2_,C_3_,D_2_,E_1_,E_2_,E_3_), 10 µm (B_1_).

Among subcortical areas, CaCyBp immunoreactive cells were localized to the amygdala (Supplementary Fig. 5C,C_1_), putamen (Supplementary Fig. 5D) and thalamus (Supplementary Fig. 5E). Unexpectedly, the substantia nigra *pars reticulata* was densely populated by CaCyBp-immunoreactive perikarya, their levels being similar to those in substantia nigra *pars compacta* (Supplementary Fig. 5F,F_1_). In the hypothalamus, CaCyBp immunoreactivity concentrated in the paraventricular (Supplementary Fig. 5G) and arcuate nuclei (Supplementary Fig. 5H). In the cerebellum, CaCyBp immunoreactivity also appeared in the molecular and internal granular layers (Supplementary Fig. 5I,I_1_), as in motor neurons of the ventral horn of the spinal cord (Supplementary Fig. 5J,J_1_).

In the adult human brain, somatodendritic CaCyBp immunoreactivity was detected in all cortical layers of the frontal and temporal lobes, many with pyramidal cell-like somatic morphology (Fig. 4D-D_2_). In the hippocampal formation, CaCyBp immunoreactivity revealed cells in the *stratum pyramidale*, with their dendrites emanating in the *strata radiatum* and *lacunosum moleculare* of the CA subfields (Fig. 4E-E_3_; Supplementary Fig. S7A-B_1_), alike cells in the hilus and granular layers of the dentate gyrus (Fig. 5E,E_1_). In subcortical regions, CaCyBp immunoreactivity dominated in the globus pallidus and caudate nucleus (Supplementary Fig. 7D,E), the thalamus, and hypothalamic nuclei (Supplementary Fig. 7F,G). In the midbrain, the substantia nigra *pars compacta* harbored remarkably dense CaCyBp immunoreactivity (Supplementary Fig. 7C,C_1_). Thus, substantial similarities remain in the regionalization, cellular density, and subcellular localization of CaCyBp immunoreactivity in adult mouse and human brains.

### CaCyBp immunoreactivity in neurons of adult mice

The identity of cells and their lineages exhibiting CaCyBp immunoreactivity is an open question. Here, we first performed multiple-labeling immunohistochemistry combining CaCyBp and NeuN to confirm that cortical and hippocampal cells with CaCyBp immunoreactivity were neurons (Fig. 5A,A_1_)^10^. To support this notion, GFAP (astrocytes)^36^, myelin basic protein (MBP; oligodendrocytes)^37^, ionized Ca^2+^-binding adapter molecule-1/allograft inflammatory factor 1 (IBA1; microglia)^38^, and nestin (neural stem cells and ependymal cells)^39,40^ did not co-localize with CaCyBp in the brain regions examined (Fig. 5B-D_1_; Supplementary Fig. 8A-A_2_).

**Figure 5.**
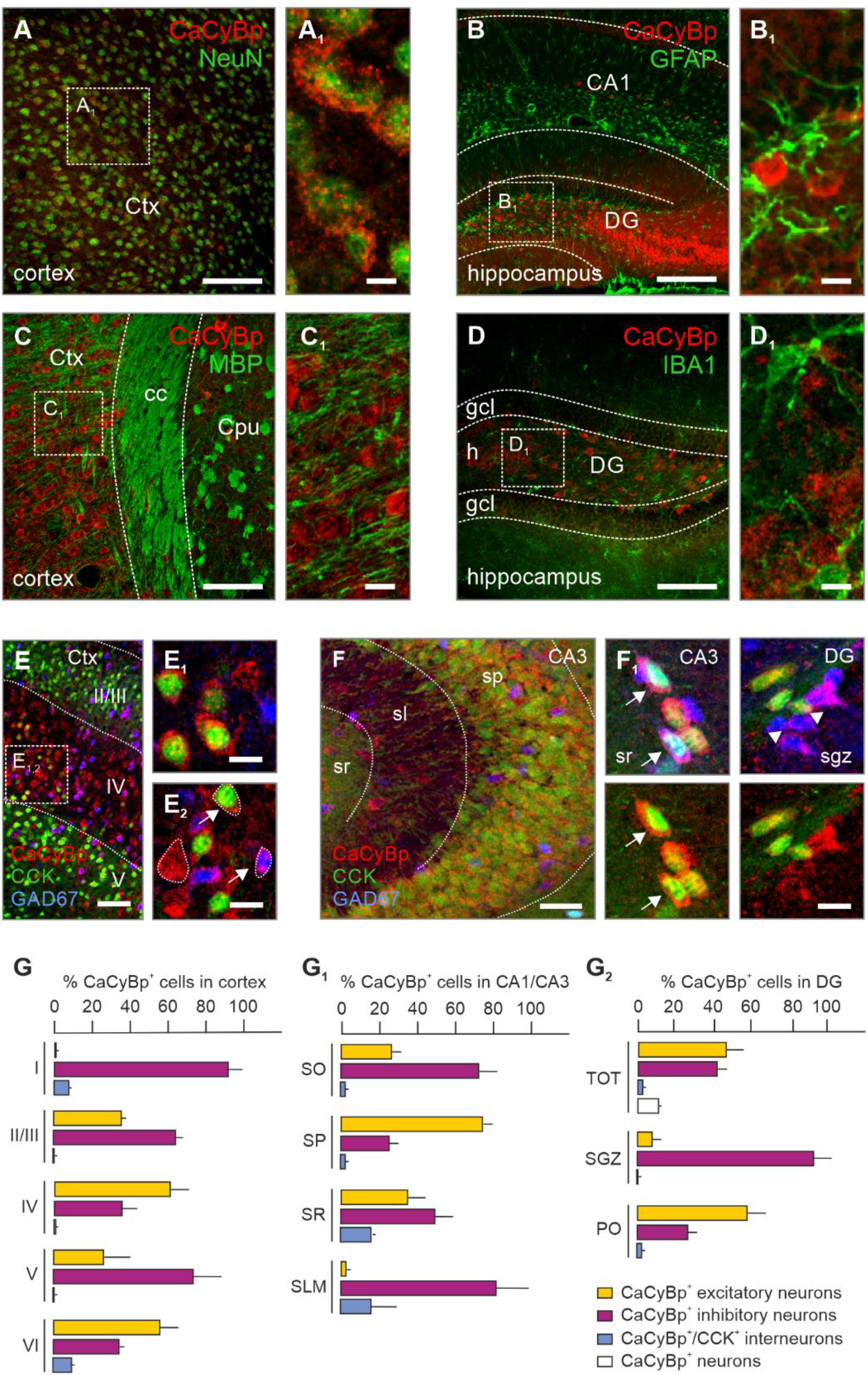
Co-localization of neuronal and non-neuronal markers with CaCyBp in the adult mouse. (**A**,**A**_**1**_) Overlap of NeuN with CaCyBp in the cerebral cortex. (**B-D**_**1**_) CaCyBp did not co-localize with any of the non-neuronal markers tested, including myelin basic protein (MBP for oligodendrocytes) in the cerebral cortex (B-B_3_), glial-fibrillary acidic protein (GFAP for astrocytes; C-C_3_) or ionized calcium-binding adaptor molecule 1 (IBA1 for microglia) in the hippocampus (D-D_3_). (**E-F**_**2**_) Immunolabeling of CaCyBp in the dual reporter mouse CCK^BAC/DsRed^::GAD67^gfp/+^ mouse, with excitatory neurons in red (DsRed alone), interneurons in green (Gfp alone) and CCK^+^ interneurons in orange (DsRed/Gfp). Arrows denote CaCyBp^+^ cells co-labelling with GAD67, while arrowheads indicate triple labelled cells. (**G-G**_**2**_) Quantification of cell numbers in the corical layers (G), hippocampal CA1-3 sub-fields (G_1_), and in the dentate gyrus (G_2_). *Abbreviations*: CA1, cornu Ammonis; cc, corpus callosum; Ctx, cortex; CPu, caudate putamen; DG, dentate gyrus; glc, granular cell layer; h, hilus; slm, stratum lacunosum moleculare; so, stratum oriens; sp, stratum pyramidale; sr, stratum radiatum;. *Scale bars* = 100 µm (E,F); 50 µm (A,B,C,D), 20 µm (A_1_,B_1_,C_1_,D_1_,E_1_,F_1_,F_2_), 10 µm (A_1_,A_2_).

We then used CCK^BAC/DsRed^::GAD67^gfp/+^ reporter mice^41^ to test if either glutamatergic neurons (DsRed only) or GABA neurons (GFP only), including their cholecystokinin (CCK)-expressing subset (dual labelled), or both, were immunoreactive for CaCyBp. In the cerebral cortex, most glutamatergic neurons with CaCyBp immunoreactivity were found in layers 4 (31.5 ± 0.1%) and 6 (37.8 ± 5.5%; Fig. 5E-E_2_,G; Supplementary Table 2). GABA interneurons with CaCyBp immunoreactivity were present in layers 4 (15.7 ± 0.2%), 5 (40.3 ± 4.3%), and 6 (18.8 ± 0.5%). CaCyBp/DsRed/GFP triple-labelled interneurons mostly populated layer 6 (95.5 ± 2.63%; Fig. 5E_2_,G)^42^.

In the hippocampus, CaCyBp immunoreactivity overlapped with DsRed in 74.5 ± 5.1% of pyramidal neurons in CA1-3 (*n* = 4; Fig. 5F,G_1_). Conversely, the majority of CaCyBp immunoreactivity in *strata radiatum* (48.8 ± 3.3%), *oriens* (71.9 ± 12.7%), and *lacunosum moleculare* (82.2 ± 3.5%) localized to GFP^+^ interneurons (Fig. 5F,G_1_; Supplementary Table 2). Triple labelling in putative CCK interneurons was also found, albeit less frequently, in *strata radiatum* (15.2 ± 2.7%), and *lacunosum moleculare* (16.3 ± 2.2%; Fig. 5F-F_2_,G_1_). In the dentate gyrus, excitatory neurons with CaCyBp immunoreactivity were mostly found in the hilus (57.5 ± 15.4%). Conversely, CaCyBp-immunopositive interneurons concentrated in the subgranular zone (92.4 ± 3.7%). CaCyBp/DsRed/GFP triple-labeled interneurons were seen in the hilus (2.5 ± 0.5%), but not in the subgranular zone (Fig. 5G_2_). In the hilus, a contingent of neurons that harbored CaCyBp immunoreactivity was negative for both GFP and DsRed (13.4 ± 1.6%), likely being excitatory mossy cells (Fig. 5G_2_). These data suggest that CaCyBp is a pleiotropic marker of adult neurons, including glutamatergic and GABAergic subtypes.

Next, we asked whether CaCyBp immunoreactivity also decorated other classes of neurons in the nervous system. For dopaminergic systems, CaCyBp immunoreactivity was detected in tyrosine hydroxylase (TH)-positive neurons of both the ventral tegmental area and substantia nigra *pars compacta* (Supplementary Fig. S8B,B_1_)^43^. For cholinergic systems, CaCyBp immunoreactivity existed in choline-acetyl-transferase (ChAT)^+^ interneurons of the dorsal striatum (Supplementary Fig. 8C,C_1_)^44^, as well as neurons in the medial septum and horizontal diagonal band of Broca (Supplementary Fig. S8D-E_2_), alike in motor neurons of the ventral spinal horn (Supplementary Fig. 8F,F_1_)^45^. For neuropeptidergic cells, CaCyBp immunoreactivity overlapped with both corticotropin releasing hormone and arginine vasopressin in the paraventricular nucleus of the hypothalamus (Supplementary Fig. S8G,G_1_)^46^.

### S100A6 expression in the adult brain

Calcyclin (S100A6) is the most studied interaction partner for CaCyBp^20^. It is therefore mandatory to assess S100A6 localization in relation to CaCyBp, at least in adult mice. S100A6-like immunoreactivity in organ systems of the mouse reminisced CaCyBp distribution, with maximum detected levels in brain, gastrointestinal tract, liver, reproductive organs, and skin (Supplementary Fig. S1B; *see also*: https://www.proteinatlas.org/ENSG00000197956-S100A6/tis-sue#expression_summary). Within the brain, S100A6 was mostly detected within or around white matter tracts in the corpus callosum (Fig. 6A-B_1_), hippocampal fimbria (Fig. 6B_2_), and internal capsule (Fig. 6B_3_). S100A6 immunoreactivity accumulated in cells with astrocyte-like morphology, but not neurons, as well as ependymocytes that lined the ventricular system (Fig. 6A,C). When using either immunohistochemistry or *in situ* hybridization, we found that S100A6 co-localized with GFAP (Fig. 6C-D_2_)^36^. These data suggest that S100A6-CaCyBp interactions could be intercellular, with astrocytes serving as a source of S100A6 for neuronal responses by CaCyBp-based ligand recognition^10,20^.

**Figure 6.**
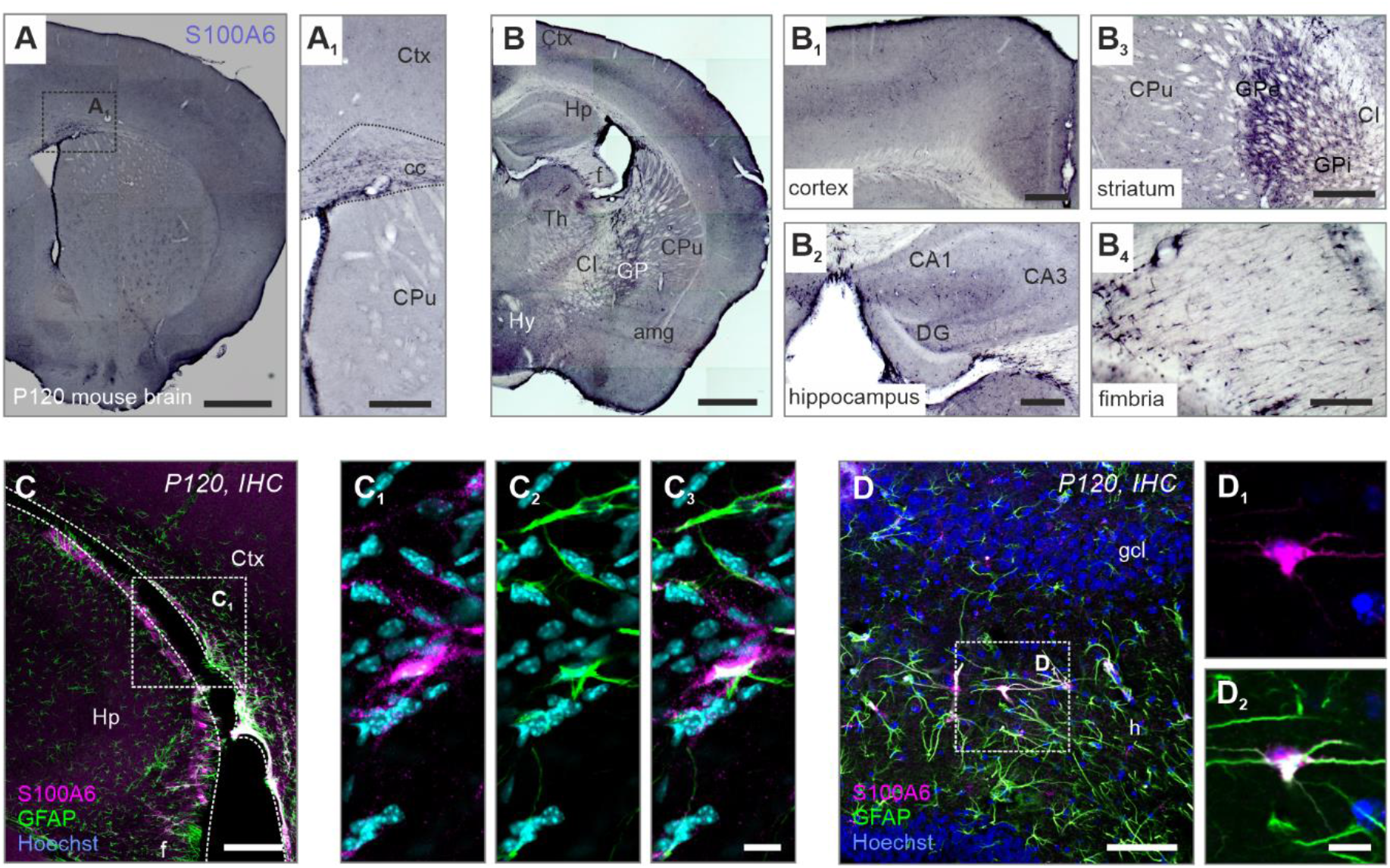
S100A6 localization in the brain. (**A-B**_**4**_) In the adult mouse forebrain, S100A6 expression was confined to astrocyte-like cells in the corpus callosum (A_1_), cortical layers (B,B_1_), hippocampus (B_2_) globus pallidus (B_3_) and fimbria (B_4_). (**C-D**_**2**_) Immunohistochemistry with the astrocyte marker GFAP confirmed the presence of S100A6 in astrocytes in limbic regions. *Abbreviations*: amg, amygdala; CA1, cornu Ammonis; CI, capsula interna; Ctx, cortex; CPu, caudate putamen; DG, dentate gyrus; gcl, granular cell layer; GPe, globus pallidus externus; GPi, globus pallidus internus; h, hilus; Hp, hippocampus; Hy, hypothalamus; Th, thalamus. *Scale bars* = 500 µm (A,B), 200 µm (C), 50 µm (A_1_,B_1_,B_2_,B_3_,B_4_,C_1_), 20 µm (D), 10 µm (D_1_).

### S100A6-CaCyBp signals affect mitochondrial protein turnover and bioenergetics

As S100A6 and CaCyBp have been implicated in mitochondrial respiration and stability in hematopoietic stem cells and immortalized cell lines^47,48^, we tested if they localized to mitochondria in neurons, too. By means of immunoelectron microscopy, we found immunogold particles inferring CaCyBp localization to mitochondrial membranes of hippocampal neurons *in vivo* (Fig. 7A,A_1_). We then used structured illumination microscopy to show that CaCyBp immunoreactivity overlapped with the OXPHOS complex, a multi-subunit protein complex (I-V) in the inner mitochondrial membrane to generate ATP via the electron transport chain^49^, particularly in the perinuclear cytoplasm of cultured hippocampal neurons (Fig. 7B-B_2_). Cellular fractionation further supported the notion that CaCyBp could be partitioned to mitochondria in the hippocampus (Fig. 7C). The finding that mitochondrial fractions from all hippocampal subfields contained CaCyBp was in line with our anatomy findings (Fig. 7C).

**Figure 7.**
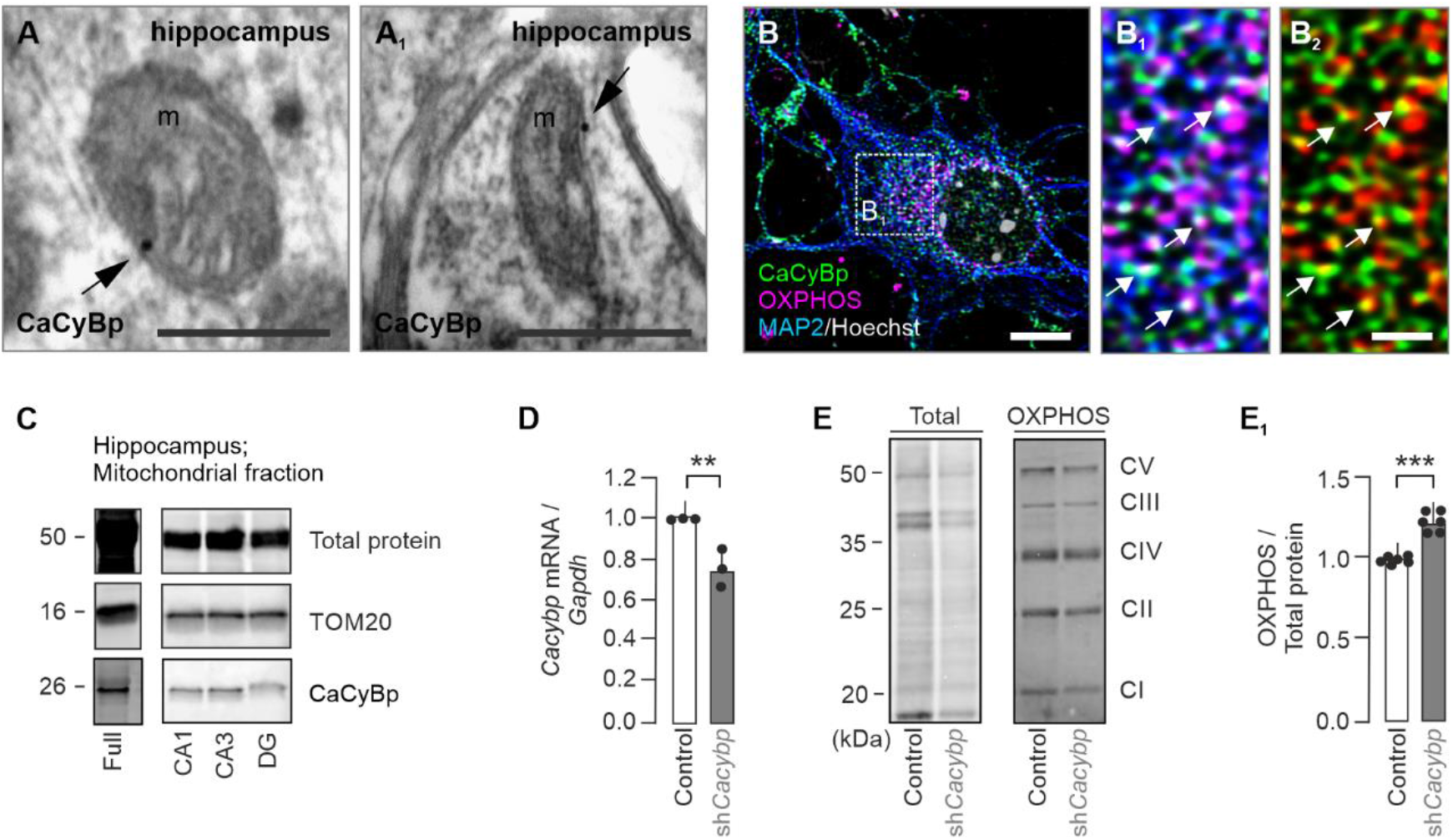
CaCyBp expression in hippocampal mitochondria. (**A**) Electron microscopy images showing CaCyBp-like immunoreactivity in mitochondria (m) in the mouse hippocampus. (**B-B**_**2**_) High-resolution laser-scanning microscopy revealed members of the OXPHOS complex and CaCyBp in close proximity in the perinuclear cytoplasm of cultured MAP2^+^ neurons. (**C**) Cellular fractionation resolved CaCyBp in mitochondrial fractions in hippocampal subfields. (**D-E**_**1**_) Treatment with a short hairpin (sh) RNAi pool against *Cacybp* mRNA reduced CaCyBp protein levels (D), while increasing members of the OXPHOS complex as compared to total protein levels (E,E_1_). Bar graphs show means ± s.e.m. \*\**p* < 0.01; \*\*\**p* < 0.001. *Scale bars* = 12 µm (B,B_1_), 500 nm (A,A_1_).

Then, loss-of-function experiments for CaCyBp were performed. Partial reduction (∼25%) of *Cacybp* expression in primary neurons by short-hairpin RNAs (shRNAs; Fig. 7D)^10^ resulted, firstly, in slowed protein turnover as per reduced protein load. Conversely, the amount of OXPHOS complex proteins disproprotionately increased (Fig. 7E,E_1_), suggesting that CaCyBp could constitutively modulate^10^ OXPHOS availability. This notion is supported by the observation that S100A6 was not detected in mitochondria of neurons under control conditions (Fig. 8A,B at’0 time point’). Yet, exposure of primary neurons to recombinant S100A6 (20 ng/ml) led to S100A6 accumulation in mitochondria from 6 h on, as judged by both immunoelectron microscopy (Fig. 8A_1_) and Western blotting of subcellular (mitochondria-enriched) fractions (Fig. 8B,B_1_). These data are compatible with S100A6 released from astrocytes and taken up by neurons (*see also* Fig. 6)^10^.

**Figure 8.**
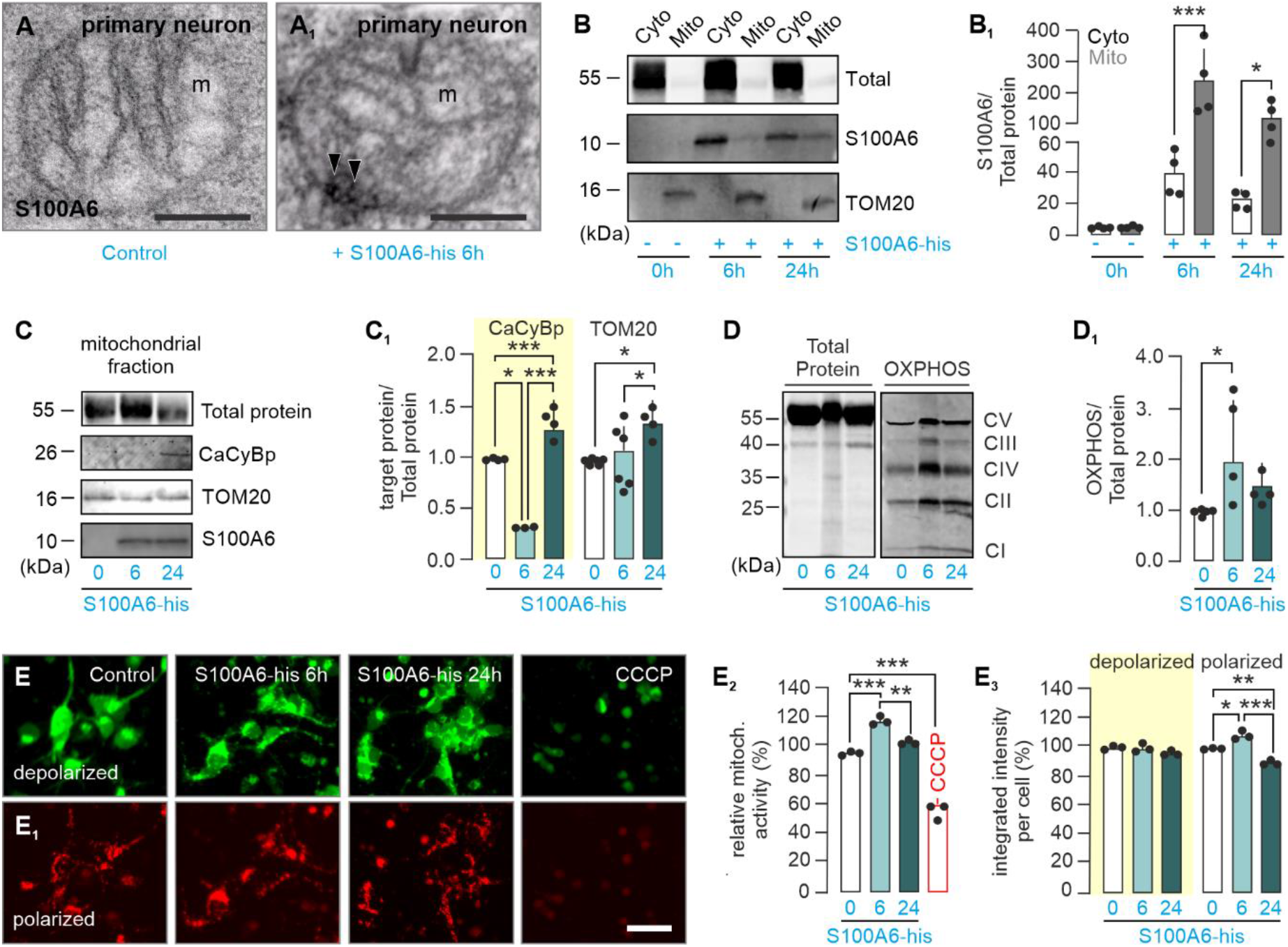
S100A6-CaCyBp affects mitochondrial protein turnover. (**A**) Electron microscopy images showing accumulation of S100A6-like immunoreactivity in mitochondria (m) of cultured primary neurons after exposure to 20 ng recombinant S100A6 for 6 hours. (**B**,**B**_**1**_) Cellular fractionation suggested the time-dependent accumulation of S100A6 in mitochondrial fractions of cultured neurons. (**C-D**_**1**_) After S100A6 exposure, CaCyBp, TOM20 (C,C_1_) and OXPHOS (D,D_1_) levels were differentially regulated. (**E-E**_**3**_) Mitochondrial membrane potential (MMP) increased after S100A6 treatment. CCCP was used as a positive control to induce a reduction in MMP. Bar graphs show means ± s.e.m. \**p* < 0.05; \*\**p* < 0.01; \*\*\**p* < 0.001. *Scale bars* = 40 µm (E_1_), 500 nm (A,A_1_).

S100A6 exposure significantly reduced CaCyBp levels in mitochondrial fractions after 6 h in neurons, with an increase at 24 h (Fig. 8C,C_1_). In harmony with data from shRNA experiments (Fig. 7E,E_1_), the lack of CaCyBp at 6 h increased mitochondrial load of both TOM20, a mitochondrial input receptor subtype^50^, as well as members of the OXPHOS complex, but not at 24 h (Fig. 8C_1_-D_1_). As the OXPHOS pathway maintains the mitochondrial membrane potential (MMP; ΔΨ*m*), which is the difference in electrical charge across the inner mitochondrial membrane that a cell uses to generate ATP, we tested whether S100A6 could indeed enhance the MMP. After 6 h, but not 24 h, of S100A6 exposure, we found significantly increased MMP levels (Fig. 8 E-E_3_). This suggest that S100A6 could inhibit CaCyBp, scale its amount in neuronal mitochondria, improve mitochondrial respiration, thus increasing ATP production10,51.

## Discussion

Our study revealed the cellular localization of CaCyBp in developing mouse and human organs, with a particular focus on the brain. While the spatiotemporal expression of CaCyBp during development and in adulthood is similar, we found variations in distribution, including transient or delayed expression: e.g., CaCyBp labelled a cohort of immature neuroblasts in the dentate neuroepithelium, a neurogenic source to granule cells by E18.5. This timing is compatible with the birth, migration, and maturation of granule cells, with the arrival of their first contingent to their final locations by P5^52,53^. From P6 into adulthood, we could not detect CaCyBp in the mouse granular cell layer, likely indicating a transient expression of CaCyBp in cells destined to this layer (similarly to classical Ca^2+^-binding proteins). In parallel, CaCyBp-containing immature calretinin-immunopositive neurons^34^ were detected in the adult subgranular zone, a remnant of the embryonic dentate neuroepithelium responsible for lifelong neurogenesis, suggesting a maintained role for developmental cellular processes in adulthood. Conversely, CaCyBp expression remained in the adult human granular cell layer as compared to the mouse, additionally emphasizing evolutionary divergence in the control of neuronal processes.

As we find CaCyBp expression in immature neurons in neurogenic niches, including the ventricular zones of the cortex and hypothalamus, as well as in migratory corridors, we posit that it tunes cellular bioenergetics for neuronal development. Our previous work demonstrated that the gliotransmission mediated by astrocyte-derived S100A6 targeting CaCyBp in neurons controls neurite outgrowth in mouse cortical neurons and its disruption can lead to altered cortical layer-patterning^10^. Additionally, in opossum cortical neurons, disrupted CaCyBp availability altered neuronal complexity, without affecting the proliferation and lineage segregation of their progenitors^54^. These actions are likely independent of S100A6 and attributed to CaCyBp as a constitutively-active regulator of transcriptions in neurons^55^. For instance, CaCyBp directly affects the availability of brain-derived neurotrophic factor (BDNF), a morphogen promoting progenitor proliferation, differentiation, and neurite outgrowth^56^, through regulating the activity of upstream ERK1/2 and cAMP response element-binding protein (CREB) signaling^57^. Alternatively, its direct regulation of β-catenin could facilitate progenitor proliferation and neuronal differentiation^58^ while its involvement in cytoskeletal instability through actin polymerization driving neurite outgrowth^8^. The importance of CaCyBp-mediated maintenance signals is also highlighted by its increased expression in embryonic cortices and altered neural progenitor proliferation in *Ts1Cje* mice modelling the neurodevelopmental disorder Down syndrome^59^. Thus, the early expression of CaCyBp during brain maturation suggests that either S100A6-dependent or constitutive signaling, or both, maintain neuronal morphogenesis.

In the adult brain, we find CaCyBp expressed by a wide variety of neurons with diverse neurotransmitter identities. For instance, CaCyBp located to both CCK^+^ excitatory pyramidal neurons, as well as multiple GABAergic interneuron subtypes (GAD67^+^ and CCK^+^/GAD67^+^) in the mouse hippocampal subfields^41^. CaCyBp was likewise detected in cholinergic neurons of the striatum, dopaminergic neurons of the ventral tegmental area and substantia nigra *pars compacta*, and co-labeled with neuropeptides in the hypothalamus. We therefore consider CaCyBp a ubiquitously expressed neuronal protein, driving omnipresent mechanisms of protein de-phosphorylation, cytoskeletal dynamics, gene expression^7^, and mitochondrial activity irrespective of neuronal identity. S100A6, the natural ligand for CaCyBP, is expressed exclusively in glial lineages, including ependymoglial cells. Therefore, S100A6 qualifies as a gliotransmitter, as being produced and secreted solely by astrocytes and acting on its target present exclusively in neurons. These characteristics also match fundamental criteria defining neurotransmitters, as outlined by John Eccles, for cell-specific chemical transmitters^60^. Thus, S100A6, together with many astrocyte-specific molecules that signal to neurons, constitute a class of true gliotransmitters mediating astroglial-neuronal communication. It is not entirely unexpected that a center for S100A6-CaCyBp signaling is in mitochondria as the CaCyBp-mediated change in the mitochondrial membrane potential is a means to affect energy production, with subsequent and substantial impact on energy-demanding protein translation, trafficking, and maturation for morphogenesis.

Above all, dysregulation of CaCyBp expression could be a biomarker^17^ to indicate negative cellular pressures and metabolic constraints. Alternatively, it could be considered an adaptive mechanism to stabilize cellular function, especially when keeping its role in mitochondrial activity and cellular stress responses in mind^9,10^. Indeed, in the aging brain, as well as in neurodegenerative disorders such as Alzheimer’s disease and frontotemporal dementia characterized by mitochondrial malfunction^61^, altered CaCyBp expression is thought to contribute to cytoskeletal instability by dephosphorylating tau protein, a culprit for neurocircuit disruption^14,62^. In a transgenic model of Huntington’s disease^26^, CaCyBp was found two-fold upregulated in the striatum, but downregulated in human Parkinson’s disease cases^64^. The latter most likely directly determines disease progression, as CaCyBp acts as a HSP90 co-chaperon whose interaction prevents α-synuclein aggregation into the characteristic Lewy bodies^65^. Global gene expression profiling flagged CaCyBp as a potential marker for patients suffering from bipolar disorder^66^, while disruptions to its interacting partner S100A6 are characteristic for sclerotic sites in amyotrophic lateral sclerosis^67^ and epileptic brains^68^, and it is found decreased in multiple brain regions during chronic mild stress in mice^69^. In sum, the ubiquitous expression of CaCyBp over many different neuronal subtypes, its interaction with many partners regulating key cellular processes and its involvement in neurodegenerative pathologies stresses an unexpectedly substantial contribution for CaCyBp in both developmental and adult neuronal functions.

## Materials and methods

### Animals and ethical approval of animal studies

Mice (C57Bl6/J) were housed in perplex cages on a 12h/12h light/dark cycle (lights on at 08:00 h) in a temperature (22 ± 2 °C) and humidity (50 ± 10%)-controlled environment. Food and water were available *ad libitum*. Embryos and tissues were obtained from timed matings with the day of vaginal plug considered as embryonic day (E) 0.5. Experiments on live animals conformed to the 2010/63/EU European Communities Council Directive and were approved by the Austrian Ministry of Women, Science and Research (66.009/0277-WF/V/3b/2017 and 2024-0.518.940). Particular effort was directed towards minimizing the number of animals used and their suffering during experiments.

### Immunohistochemistry, microscopy, and imaging

Adult male mice were transcardially perfused with ice-cold 0.1 M phosphate buffer (PB; 20 ml), followed by 4% paraformaldehyde (PFA) in PB (100 ml at 3 mL/min flow speed). Postnatal (P)120 adult brains, embryonic (E)18.5 whole embryo heads (or their dissected brains) were (post-)fixed by immersion in 4% PFA overnight. After equilibrating in 30% sucrose for 48–72 h, coronal serial sections (1-in-6 series) were cryosectioned (Leica CM1850) at a thickness of 50 μm (adult brain), 25 μm (whole embryo) or 20 μm (embryonic brain). Spinal cord (SC) was dissected out and post-fixed in the same fixative at 4 °C for 90 min, followed by immersion in 10% (vol/vol%) sucrose in 0.1 M PB containing 0.01% sodium azide (Merck) and 0.02% bacitracin (Sigma). Tissues were kept in 10% sucrose solution for cryoprotection at 4 °C for 2 days. Tissues were then trimmed and embedded in optimal cutting temperature compound (OCT; HistoLab AB), snap-frozen in liquid carbon dioxide, and sectioned on a cryostat (ThermoFisher NX70) at a thickness of 20 µm. Sections were routinely thaw-mounted onto SuperFrost Plus^+^ glass slides. Free-floating and glass-mounted sections were processed as described^70^ for either fluorescence or chromogenic light microscopy (using 3,3’-diaminobenzidine (DAB) with 0.1% H_2_O_2_ as substrate).

For fluorescence histochemistry, free-floating sections from *n* = 4 male adult mice were extensively rinsed in PB, and blocked with PB containing 5% normal donkey serum (NDS; Jackson ImmunoResearch; #017000121), 1% bovine serum albumin (BSA; Sigma; #017000121), and 0.3% Triton X-100 (Sigma) at 21-24 °C for 2 h to reduce non-specific labeling. Next, sections were incubated with a combination of primary antibodies diluted in PB also containing 2% NDS, 0.1% BSA and 0.3% Triton X-100 at 4 °C for 72 h (Supplementary Table 1). DyLight Fluor 488, 560, or 633-tagged secondary antibodies (1:300; Jackson ImmunoResearch) were used for signal detection. Hoechst 33,342 (Sigma) or 4′,6-diamin-2-phenylindol (DAPI, Sigma; #D8417) were routinely used as nuclear counterstains. After extensive washing in PB and a final rinse in distilled water, free-floating sections were mounted onto fluorescence-free glass slides and coverslipped with Entellan (in toluene, Merck). Glass-mounted embryonic sections of both sexes were processed in fashion identical to above, except that either Aquamount (Dako) or Mowiol 4-88 (Roth) containing 1,4-diazabicyclo[2.2.2]octane (DABCO, Sigma) was used for coverslipping. Images were acquired on a Zeiss 880LSM confocal laser-scanning microscope equipped for maximal signal separation and spectral scanning/unmixing.

For chromogenic histochemistry, sections were washed with PB (3×, 5 min each), and then treated with 1% H_2_O_2_ for 10 min to block endogenous peroxidases. After rinsing, sections were exposed to a blocking solution (*see above*) at 21-24 °C for 2 h. Sections were then incubated with a rabbit anti-CaCyBp primary antibody or with a rabbit anti-S100A6 primary antibody (Supplementary Table 1) for 72 h. After washing (3×), sections were exposed to biotinylated anti-rabbit secondary antibodies (1:200; Jackson ImmunoResearch, 20–22 °C, 2 h), rinsed again in PB, and then exposed to preformed avidin-biotin-peroxidase complexes (1:200, ABC Elite; Vector Laboratories) at 4 °C overnight. Next, biotin-labeled cells were revealed using a peroxidase reaction with DAB alone (5 mg/ml, 0.03% H_2_O_2_ as substrate, brown reaction product) or in combination with nickel–ammonium sulphate using the glucose oxidase method for H_2_O_2_ generation (blue particulate products)^71^. Sections were subsequently mounted onto clean slides, dried, dehydrated in an ascending alcohol gradient (30%, 50%, 70%, 100% ethanol, 10 min each), cleared in xylene for 20 min and coverslipped with DPX mounting medium (Sigma). Tissues were inspected on a Zeiss Apotome.2 microscope equipped with an Axiocam 506 camera.

### Electron microscopy

Electron microscopy was performed as published^72^. Briefly, after fixation, tissues were rinsed in PB (3×) and then incubated in 0.1 M Tris-NaCl buffer supplemented with 0.1% Tween-20 (TNT), 1% BSA and 1% glycine for 30 min. This was followed by exposure to anti-CaCyBp or anti-S100A6 antibody at 4°C overnight (Supplementary Table 1). Cells were then rinsed in the above buffer and incubated with goat anti-rabbit F(ab’)2 coupled to 0.8 nm gold particles (1:100; Aurion) in PB-TNT-BSA for 2 h at 22–24°C. After fixation with 1% glutaraldehyde (10 min) and rinsing in water, gold particles were silver-enhanced using the HQ Silver Enhancement Kit (Nanoprobes; 5 min at 22– 24°C) before post-fixation in 1% OsO4 for 1 h. Next, cells were embedded (Epoxy resin), and after polymerisation, regions of interest were identified under a stereomicroscope, isolated with a razor blade and re-embedded in Epon (Fisher Scientific). Specimens were observed by using a JEOL electron microscope.

### Human tissue collection and immunohistochemistry

Adult (57-63 years) and fetal (gestational week (GW) 27-34) brains (*n* = 3 samples/age; mixed for sex) were collected at the Neurobiobank of the Institute of Neurology, Medical University of Vienna, Austria^31,32^. Tissues were obtained and used in compliance with the Declaration of Helsinki, and following institutional guidelines approved by the Ethical Committee of the Medical University of Vienna (No.104/2009 and No. 396/2011 for fetal and adult brains samples, respectively). Fetal brain tissue was obtained from spontaneous or medically-induced abortions. Only brains of fetuses (15–34 GW), and adults (50–63 years) whose cause of death was unrelated to genetic disorders, head injury, neurological diseases or other known diseases (e.g., infections) were included. Tissues were kept at 4 °C for 24–48 h due to local regulations before immersion fixation in saturated formalin (10%) on average for 34–37 days before embedding in paraffin. Three-μm sections of formalin fixed, paraffin-embedded tissue blocks were mounted on pre-coated glass slides (StarFrost) and stained as per published protocols^31,32^. Following deparaffinization and rehydration, sections were incubated in a solution of H_2_O_2_ (3%) in absolute methanol for 10 min to block endogenous peroxidases. After washing with distilled water, sections were pre-incubated in low pH EnVision FLEX antigen retrieval solution (PTLink; Dako) at 98 °C for 20 min, exposed to fetal calf serum (10%) for 10 min, and subsequently stained with rabbit anti-CaCyBp antibody (Supplementary Table 1) at 4 °C for 72 h. The EnVision detection kit (K5007, ready-to-use, DAKO) was used to visualize the horseradish peroxidase/DAB reaction product with H_2_O_2_ substrate (0.01%). Sections were counterstained with hematoxylin-eosin, dehydrated in an ascending series of ethanol, cleared with xylene, covered with Consul-Mount (Thermo Scientific), and viewed under a Nikon Eclipse E400 microscope.

### *In situ* hybridization (HCR-FISH)

Fresh-frozen E18.5 mouse brains were cryosectioned at a thickness of 16 μm and collected on SuperFrost^+^ glass slides (ThermoFisher). Sections were immersed in 4% PFA solution for 15 min, and dehydrated in an ascending ethanol gradient (50%, 70%, and 100%; 5 min each). The HCR RNA-FISH protocol for ‘*fresh frozen or fixed frozen tissue sections*’ (Molecular Instruments) was used with pre-designed *Cacybp, S100a6* and *Rbfox3* probes^10^. Samples were imaged on an LSM 880 confocal microscope (Zeiss) at 40× primary magnifications.

### RNA isolation and quantitative PCR

The RNeasy mini kit (Qiagen) was used to extract RNA from mouse cerebral tissues. DNase I was used to eliminate traces of genomic DNA. Total RNA was then reverse transcribed to cDNA in a reaction mixture using a high-capacity cDNA reverse transcription kit (Applied Biosystems). cDNA was then used for quantitative real-time PCR (CFX-connect, Bio-Rad) with pairs of PCR primers specific for mouse *Cacybp* designed with Primer Bank and the NCBI Primer Blast software (*forward*: GGT TGC TCC TCT TAC AAC AGG; *reverse*: TGA CCT CTC TGT GAA GTG CA). Quantitative analysis of gene expression was performed with SYBR Green Master Mix Kit (Life Technologies). Expression levels were normalized to glyceraldehyde-3-phosphate dehydrogenase (*Gapdh*; *forward*: AAC TTT GGC ATT GTG GAA GG; *reverse*: ACA CAT TGG GGG TAG GAA CA), a housekeeping gene, for every sample in parallel assays (*n* ≥ 3 biological replicates/group).

### Total protein determination and quantitative Western blotting

Total protein was extracted from cerebral tissues by using a modified radioimmunoprecipitation assay buffer containing 50 mM Tris-base (pH 7.4; Sigma), 1 mM EDTA (Merck), 150 mM NaCl (Fisher Chemical), 5 mM NaF (sigma,), 5 mM Na_3_VO_4_ (Sigma), 0.1% *N*-octyl-β-d-glucopyranoside (Sigma), 0.5% Triton X-100, and a cocktail of protease inhibitors (Complete, EDTA-Free; Roche). Samples were homogenized by sonication and protein lysates were pelleted at 10,000 *g* at 4 °C for 10 min. Protein concentrations were determined by Bradford’s colorimetric method to aid equal loading in Western blot experiments^73^.

To fractionate mitochondria and cytoplasm, samples were homogenized in a buffer containing 215 mM mannitol, 75 mM sucrose (Sigma); 0.1 % BSA, 20 mM Hepes (pH 7.4), 1 mM EGTA (all from Sigma), and a cocktail of protease inhibitors (Roche), and centrifuged twice at 600 *g* at 4 °C for 5 min. The supernatant was then centrifuged at 10,000 *g* at 4 °C for 10 min. The final pellet was then reconstituted as the mitochondrial fraction, whereas the supernatant was saved as the cytosol fraction. Protein concentrations were determined by the Pierce BCA Protein Assay kit. Anti-TOM20 antibodies (Supplementary Table 1) were used to verify enrichment of the mitochondria fraction.

Total protein labeling was initiated by carbocyanine (Cy)5 dye reagent (GE Healthcare) that had been pre-diluted (1:10) in ultrapure water. Samples were mixed and incubated at 21-24 °C for 5 min. The labeling reaction was terminated by adding Amersham WB loading buffer (GE Healthcare; 20 μl/sample) containing 40 μM dithiothreitol (Sigma, #10197777001). Equal amounts of the samples (20 μg/40 μl) were then boiled at 95 °C for 3 min, and subsequently loaded onto an Amersham WB gel card (13.5%). Electrophoresis (600 V, 42 min) and protein transfer onto PVDF membranes (100 V, 30 min) were at default settings in an integrated Amersham WB system (GE Healthcare) for quantitative SDS-PAGE and Western blotting of proteins with fluorescence detection. After blocking, membranes were incubated with primary antibodies overnight (Supplementary Table 1). Antibody binding was detected by using species-specific Cy3-labeled secondary antibodies (1:1,000; GE Healthcare). Membranes were dried before scanning at 560 nm (Cy3) and 630 nm (Cy5) excitation. Automated image analysis was performed with the Amersham WB evaluation software with linear image optimization, if necessary.

### Neuronal cultures, shRNA knockdown, and treatment with recombinant S100A6

Cerebral tissues of mouse embryos (mixed for sex) were isolated on either E18.5 or P0. Cells were mechanically dissociated into a single-cell suspension by papain digestion (20 U/ml, pH 7.4, 45 min; Worthington) and plated onto poly-D-lysine-coated (Sigma) 6-well (200,000 cells/well), 24-well (50,000 cells/well) or 96-well plates (30,000 cells/well), and 6-cm dishes (1x10^6^ cells/well). Cells were maintained in Neurobasal A medium (Fisher Scientific) supplemented with B27 (1%; Thermo Fisher), GlutaMAX (1%; Thermo Fisher), and penicillin/streptomycin (1%; Life Technologies). Media were replaced every other day.

shRNA knock-down of *Cacybp* was performed as previously reported^10^. Briefly, P0 cortical neurons (200,000 cells/well) were seeded in 6-well plates, grown for 48 h, then transduced with either 1 μl of adeno-associated virus, serotype 2 (AVV-2) particles (6.09 x 10^12^ genome copies/ml; GeneCopoeia; #AA02-CS-MSE091282-AVE001-01-200) to carry shRNA or equal amounts of AVV-2s to deliver scrambled controls (GeneCopoeia; #AA02-CS-CSECTR001-AVE001-01-200). Media were replaced with full Neurobasal A medium (as above) free of AAV-2 particles after 48 h, and changed every other day. Cells were then lysed for subcellular fractionation, Western blotting, and qPCR after 15 days *in vitro* (DIV).

To test the partitioning of CaCyBp to mitochondria, E18.5 neurons were plated in 24-well plates (50,000 cells/well), and allowed to grow until 4 DIV. Neurons were then immersion-fixed in 4% PFA (in 0.1 M phosphate-buffered saline [PBS]) for 1 h, and then exposed to a blocking solution consisting of 10% NDS, 5% BSA, and 0.3% Triton X-100 in PBS (pH 7.4) at 21-24 °C for 2 h. To visualize mitochondria, cultured neurons were incubated with an anti-OXPHOS antibody (Supplementary Table 1). Hoechst 33,342 (Sigma) was routinely used as nuclear counterstain, and microtubule-associated protein 2 (MAP2)^74^ to label the somatodendritic compartment of neurons *in vitro*. Images were acquired on a Zeiss Lattice SIM3 super-resolution microscope with maximal signal separation and spectral scanning/unmixing and SIM^2^ post-processing.

For the subcellular distribution and localization of S100A6, E18.5 neurons were plated at a density of 50,000 cells/well in 24-well plates or 1x10^6^ cells in 6-cm Petri dishes. After 3 DIV, neurons were exposed to 20 ng of recombinant His-tagged S100A6 protein (Abcam) for 6 and 24h and then processed for either immunocytochemistry (6 h) or subcellular fractionation (6 and 24 h; *see below*). S100A6 on mitochondria were detected by both electron microscopy and Western blotting.

### Mito-ID

E18.5 neurons were plated at a density of 30,000 cells per well in poly-D-lysine–coated 96-well plates. S100A6 treatment started on 3 DIV as indicated. The mitochondrial membrane potential (MMP) was determined by the Mito-ID Membrane Potential Detection Kit (Enzo Life Sciences). All reagents and buffers were according to the manufacturer’s protocol. After exposure to recombinant 20 ng S100A6 (Abcam) for either 6 or 24 h, neurons were washed twice with ‘Mito-ID assay buffer’. Subsequently, ‘Mito-ID MP detection reagent’ was added for 15 min. Measurements were then performed in ‘assay buffer’ in a humidified atmosphere (5% CO_2_) at 37 °C. Carbonyl cyanide m-chlorophenyl hydrazone (CCCP, 5 μM) was used as a positive control for mitochondrial depolarization. Energized mitochondria with a high MMP were detected using a rhodamine filter (excitation: 540 nm, emission: 570 nm). Depolarized mitochondria with low MMP were visualized with a fluorescein filter (excitation: 485 nm, emission: 530 nm). Experiments were performed in quadruplicate. Signals were measured separately per channel. Subsequently, the ratio of high and low MMP signals was calculated.

### Statistical analysis and figure design

The number of independent samples is indicated in either the individual figure panels or their associated legends. Likewise, the number of animals was given where relevant. All values represent the means ± s.e.m. of independent experiments. Biological variation among the samples was similar throughout. Statistical significance was determined by Student’s *t*-test (for independent groups, two-tailed) or one-way analysis of variance (ANOVA) followed by Bonferroni’s *post-hoc* correction (for multiple groups). Statistical analysis was performed in Prism 8.0 (GraphPad Software Inc.). Multi-panel images were assembled in CorelDraw v.24.0 with linear color, brightness or contrast enhancement to increase visual clarity, as necessary and appropriate.

## Acknowledgments

The authors thank Jan Mulder and Mathias Uhlén (Karolinska Institutet) for anti-S100A6 and anti-CaCyBp antibodies, Gabor G. Kovacs (Medical University of Vienna) for human tissues, G. Szabo and F. Erdelyi (HUN-REN Institute of Experimental Medicine) for reporter mice, and Alessa Reinthaler for her expert laboratory assistance. This work was supported by funding from the Austrian Science Fund FWF (10.55776/COE16, T.H.), Vetenskapsrådet (2023-03058, T.H.), the Novo Nordisk Foundation (NNF23OC0084476, T.H.), and the European Research Council (FOODFORLIFE, ERC-2020-AdG-101021016, T.H.).

## Notes

**Conflict of interest:** The authors declare no conflict of interest.

### Competing Interest Statement

The authors have declared no competing interest.

### Summary of Updates

The title has been updated, and minor spelling errors have been corrected.

