## Supplementary Information for "S100A6 gliotransmission modulates neuronal proteostasis and energy production in the mouse and human brain"

**Contents:**

- Supplementary Table 1-2
- Supplementary Figures 1-8
- Captions and Legends to Supplementary Table and Figures
- References

| Antibody | Host species | Concentration | Manufacturer |
|---|---|---|---|
| AVP | Goat | 1:50 (IHC) | Santa Cruz Biotechnology |
| CaCyBp | Rabbit | 1:1,000 (IHC, WB) | HPA #HPA025753 |
| Calbindin D28k | Mouse | 1:1,000 (IHC) | Swant #300 |
| Calretinin | Mouse | 1:1,000 (IHC) | Swant #CG1 |
| ChAT | Goat | 1:200 (IHC) | Millipore #AB144P |
| CPCA-mCherry | Chicken | 1:1,000 (IHC) | EnCor #CPCA-mCherry |
| Doublecortin | Guinea pig | 1:1,000 (IHC) | Millipore #AB2253 |
| GFP-FITC | Goat | 1:1,000 (IHC) | Abcam #Ab6662 |
| GFAP | Rabbit | 1:1,000 (IHC) | Synaptic Systems #173002 |
| Hoechst 33,342 | - | 1:10,000 (IHC) | Sigma #14533 |
| IBA1 | Goat | 1:500 (IHC) | Abcam #Ab5076 |
| MAP2 | Guinea Pig | 1:1,000 (ICC) | Synaptic Systems #188004 |
| Nestin | Mouse | 1:200 (IHC) | Millipore #MAB353 |
| NeuN | mouse | 1:1,000 (IHC) | Millipore #MAB377 |
| MBP | Mouse | 1:200 (IHC) | Boehringer Mannheim |
| OXPHOS | Mouse | 1:500 (ICC, WB) | Abcam #45-8099 |
| Parvalbumin | Guinea Pig | 1:500 (IHC) | Synaptic Systems #195308 |
| S100A6 | Rabbit | 1:1,000 (IHC, WB) | HPA #HPA007575 |
| TH | Rabbit | 1:500 (IHC) | Millipore #AB152 |
| TOM20 | Rabbit | 1:500 (ICC, WB) | Santa Cruz Biotechnology #FL-145 |
| VGLUT2 | Guinea Pig | 1:100 (IHC) | M. Watanabe (Private)[1] |
| Vimentin | Chicken | 1:500 (IHC) | Synaptic Systems #172006 |

**Supplementary Table 1. Antibodies used for immunofluorescence histochemistry and Western blotting.** Application-specific dilutions, hosts, and vendors for the antibodies specified were stated.

| Brain regions | | Excitatory neurons | Inhibitory neurons | Triple labelling | CaCyBp |
|---|---|---|---|---|---|
| | | CaCyBp + $CCK^{BAC/DsRed}$ | CaCyBp + $GAD67^{gfp/+}$ | CaCyBp + DsRed/GFP | |
| Cortex | Layer I | n.d. | 1.6 ± 0.2 | 2.3 ± 1.3 | n.d. |
| | Layer II/III | 15.1 ± 1.6 | 26.3 ± 5.2 | 2.2 ± 1.3 | n.d. |
| | Layer IV | 31.5 ± 0.1 | 15.7 ± 0.2 | n.d. | n.d. |
| | Layer V | 15.6 ± 3.8 | 40.3 ± 4.3 | n.d. | n.d. |
| | Layer VI | 37.8 ± 5.5 | 18.8 ± 0.5 | 95.5 ± 2.6 | n.d. |
| CA1/CA2 | SO | 26.3 ± 3.9 | 71.9 ± 12.7 | 1.8 ± 1.3 | n.d. |
| | SP | 74.5 ± 9.9 | 25.2 ± 15.6 | 0.3 ± 0.7 | n.d. |
| | SR | 36 ± 7 | 48.8 ± 3.3 | 15.2 ± 2.7 | n.d. |
| | SLM | 1.6 ± 0.3 | 82.2 ± 3.5 | 16.3 ± 2.2 | n.d. |
| DG | TOT | 46.2 ± 4.4 | 41.6 ± 3 | 1.9 ± 0.7 | 10.3 ± 1.6 |
| | SGZ | 7.6 ± 1.1 | 92.4 ±3.7 | n.d. | n.d. |
| | PO | 57.5 ± 15.4 | 26.7 ± 4 | 2.5 ± 0.5 | 13.4 ± 1.6 |

**Supplementary Table 2. Quantification of CaCyBp-immunoreactive cells in the mouse somatosensory cortex and hippocampus.** Quantification of $CaCyBp^{+}$ cell number in a dual reporter $CCK^{BAC/DsRed}$::$GAD67^{gfp/+}$ mouse strains at cortical layers, hippocampal CA1-2 subfields, and in the dentate gyrus (DG).

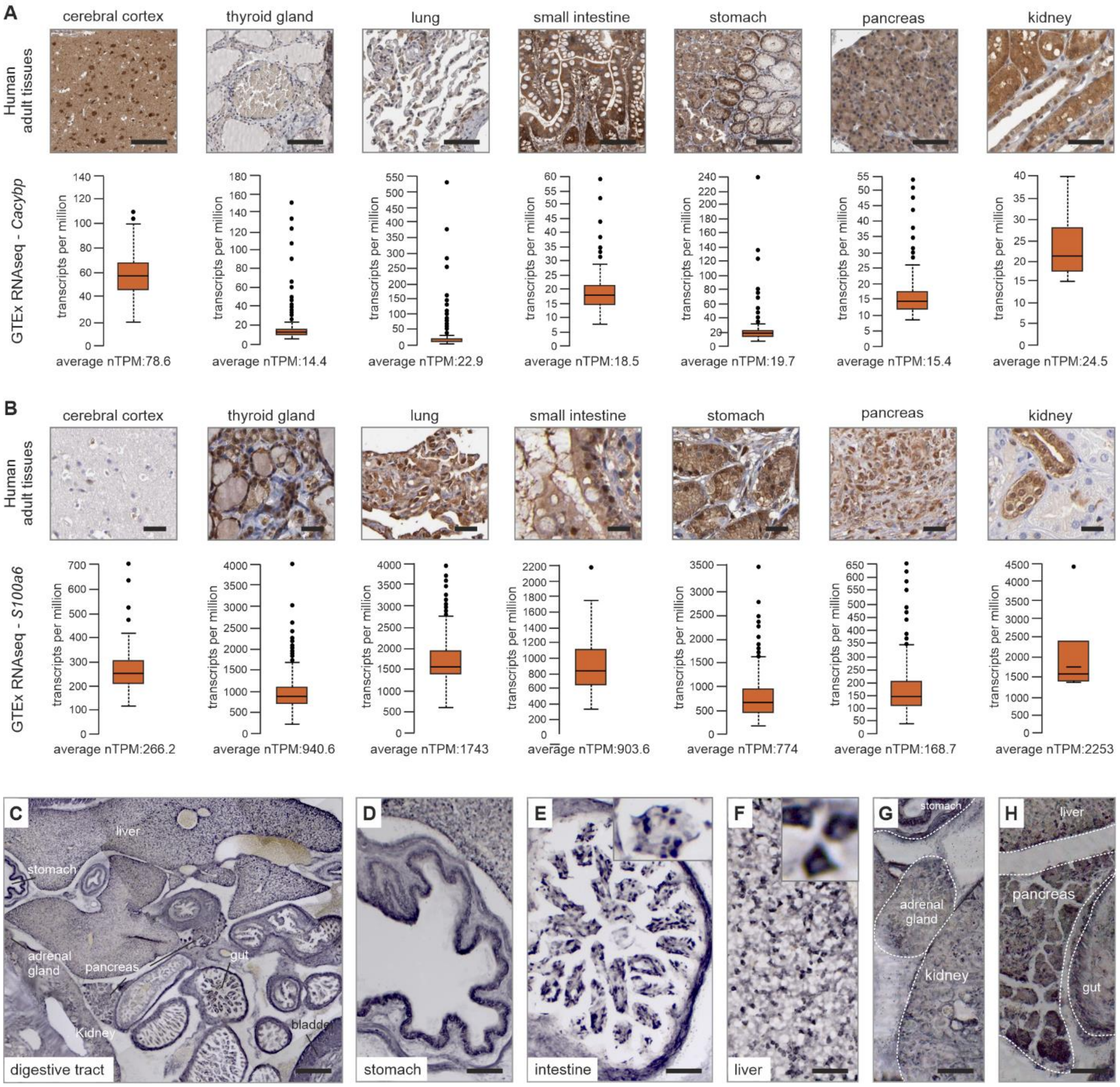


**Supplementary Figure 1. Body-wide CaCyBp and S100A6 expression profiles.** (**A,B**) Histochemical localization of CaCyBp (#HPA025753) and S100A6 protein (#HPA007575) and Genotype-Tissue Expression (GTEx) RNA-seq transcript analysis in a selection of human tissues. (**C-H**) Nickel enhancement (blue) revealed CaCyBp along the embryonic mouse gastrointestinal tract (C), with higher resolution images showing the stomach (D), intestines (E), liver (F), kidney (G), and pancreas (H). Note the predominant somatic accumulation of CaCyBp (E,F). *Scale bars* = 500 µm (C); 100 µm (A,D,E,F,G,H).

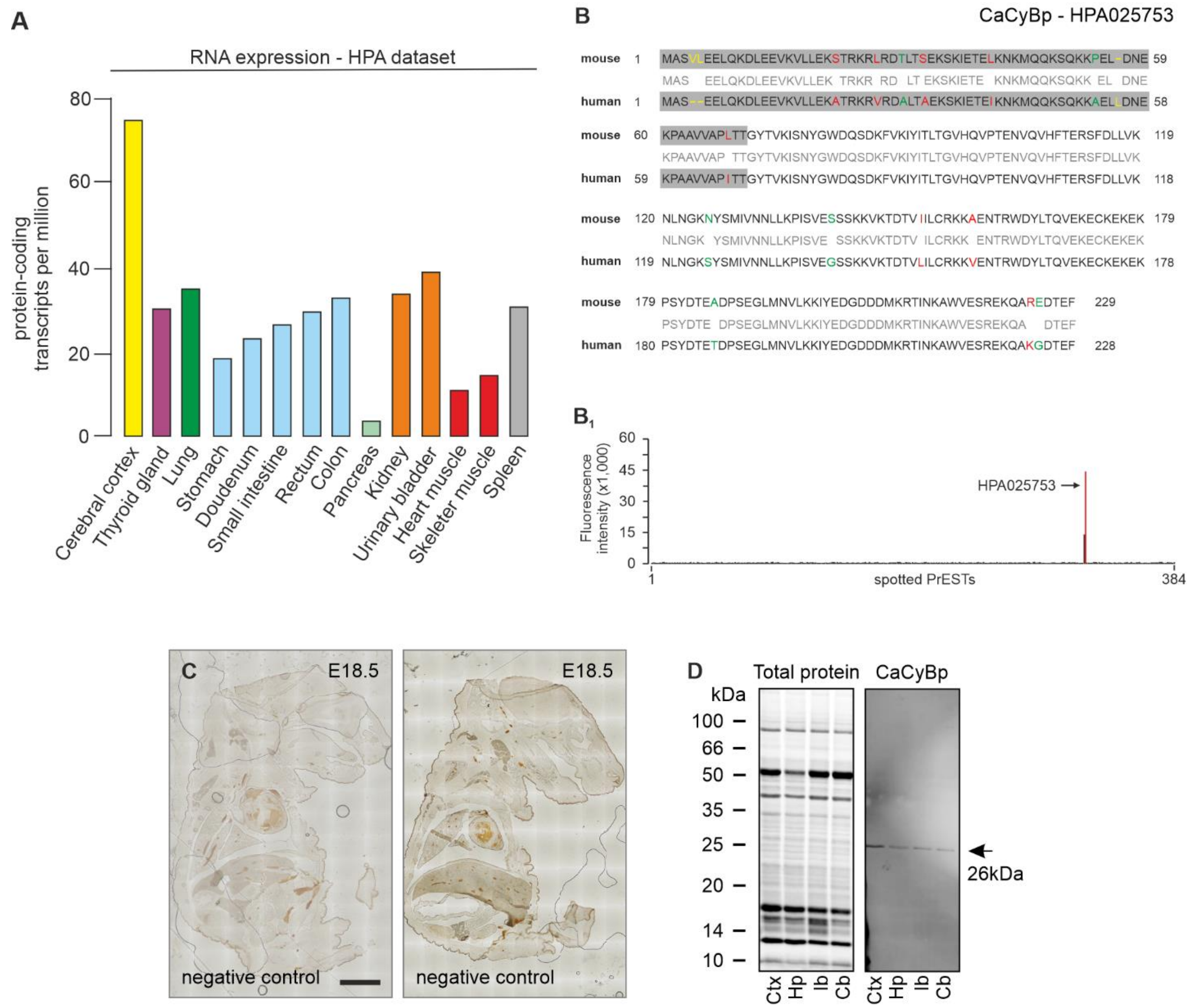


**Supplementary Figure 2. CaCyBp antibody verification.** (**A**) Expression of CaCyBP mRNA in human organ systems (obtained from the Human Protein Atlas; www.proteinatlas.org). (**B**) Homology comparison between mouse and human CaCyBp. Conserved amino acid residues are shown in red, non-conserved in green, and yellow denotes the absence of residues. Grey background highlights the antigen sequence used for antibody generation (HPA025753). (**B**$_{1}$) A protein array of 384 protein epitope signature tags (PrESTs) was used for the initial screening of antibody specificity. Note that the CaCyBp antibody used in this study only detected the CaCyBp PrEST sequence in the antigen array (*red peak*). (**C**) Immunohistochemistry with DAB amplification on full E18.5 fetuses without primary antibody incubation reveals limited background labelling. (**D**) Western blot analysis shows one band at the expected molecular weight of CaCyBp. *Abbreviations*: Cb, cerebellum; Ctx, cortex; Hp, hippocampus; Ib, interbrain (thalamus/hypothalamus). *Scale bars* = 1 mm (C).

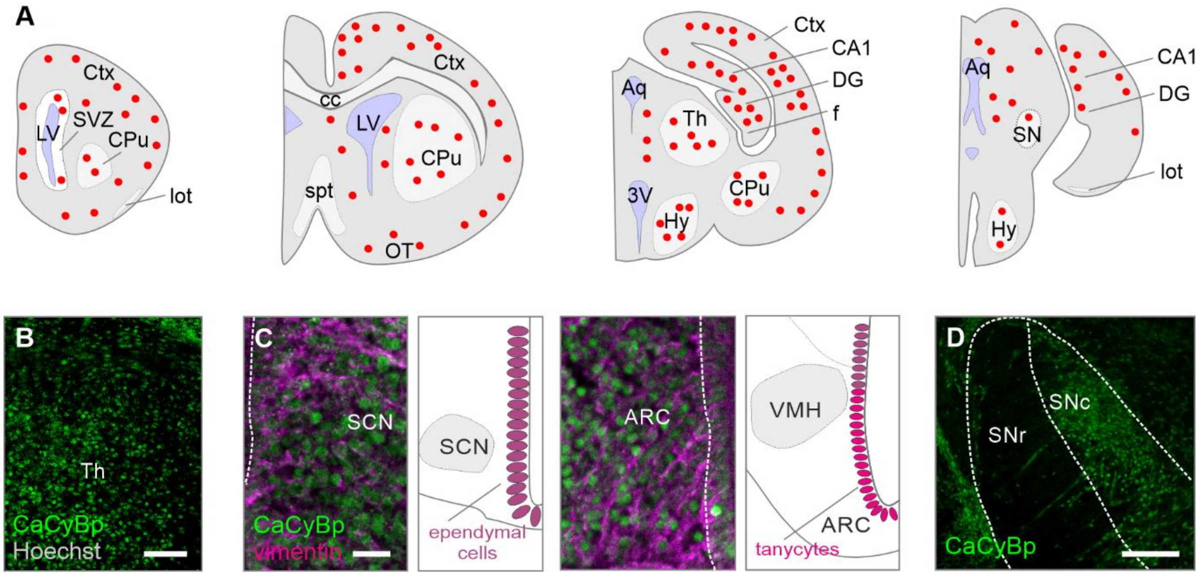


**Supplementary Figure 3. CaCyBp labelling in the fetal mouse brain.** (**A**) Mapping of CaCyBp in the E18.5 embryonic mouse brain. Red solid circles denote the relative density of CaCyBp-positive neurons in forebrain regions. (**B-D**) CaCyBp expression in the thalamus (B), hypothalamus (C) and substantia nigra (D). *Abbreviations*: 3V, third ventricle; ARC, arcuate nucleus; Aq, aquaduct; cc, corpus callosum; CPu, caudate putamen; Ctx, cortex; DG, dentate gyrus; f, fimbria; Hy, hypothalamus; LV, lateral ventricle; lot, lateral olfactory tract; OT, olfactory tract; SCN, suprachiasmatic nucleus; SNc, substantia nigra pars compacta; SNr, substantia nigra pars reticulata; spt, septum; SVZ, subventricular zone; Th, thalamus; VMH, ventromedial hypothalamus. *Scale bars* = 200 μm (B), 50 μm (C,D).

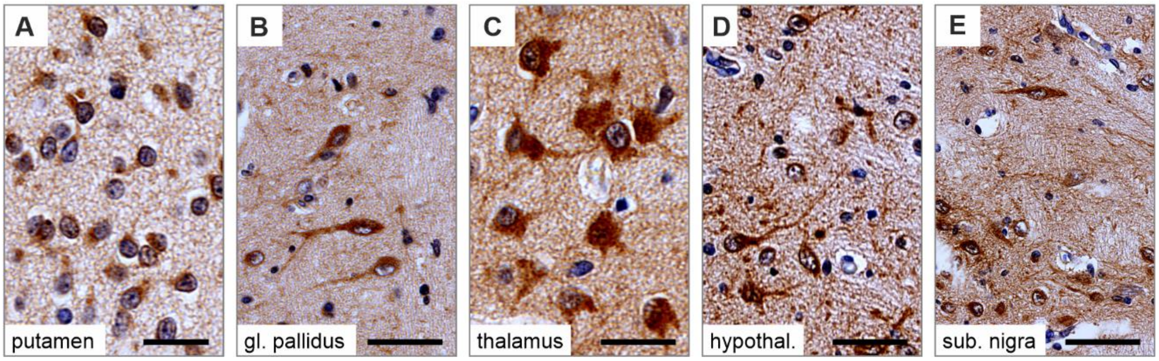


**Supplementary Figure 4. CaCyBp labelling in the fetal human brain.** (**A-E**) CaCyBp expression in the globus pallidus (A), putamen (B), thalamus (C), hypothalamus (D) and substantia nigra (E). *Scale bars* = 10 µm (A,B,C,D,E).

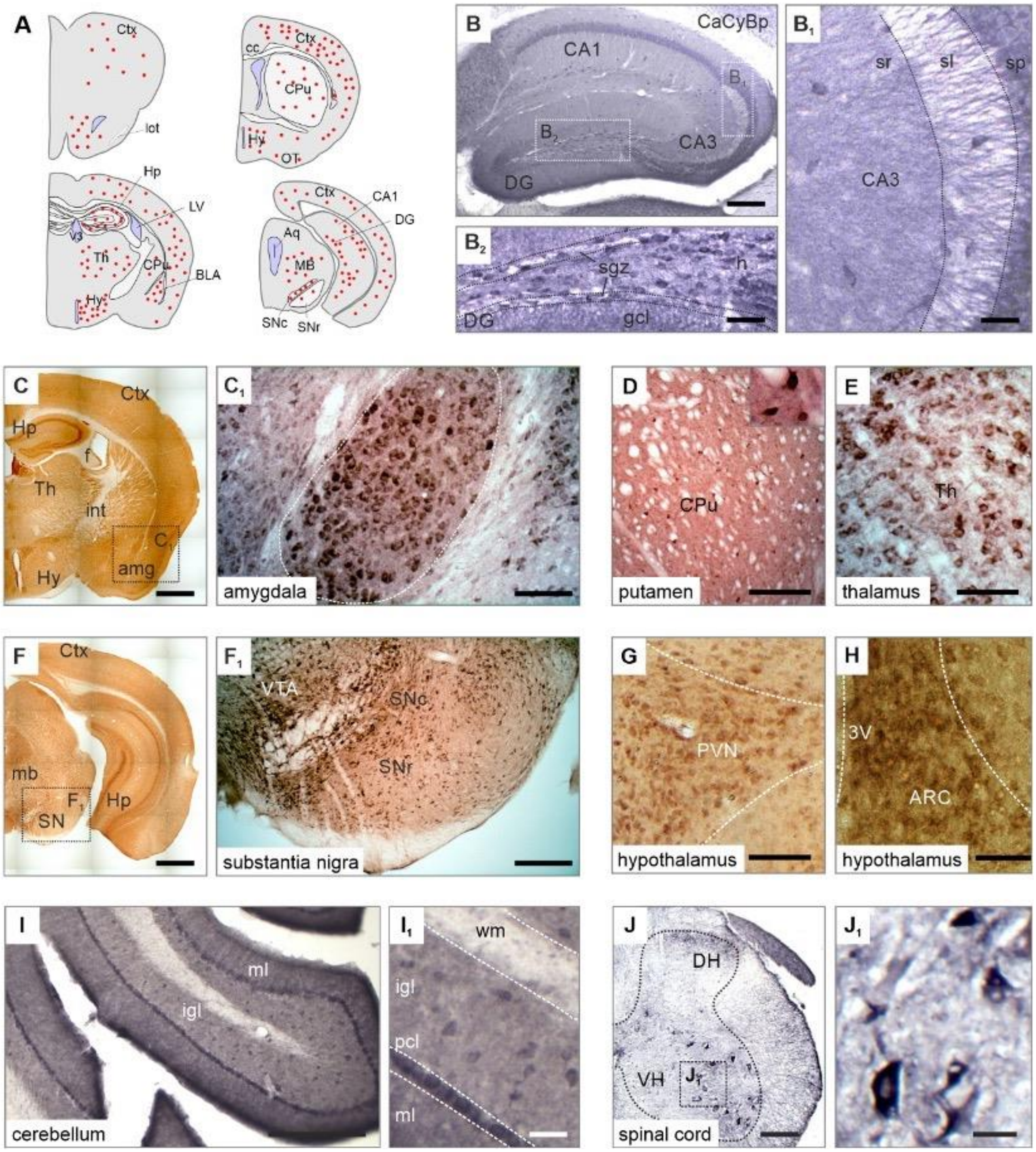


**Supplementary Figure 5. CaCyBp labelling in the adult mouse brain.** (**A**) Mapping of CaCyBp in the adult mouse brain. (**B-J$_1$**) Immunohistochemistry for CaCyBP revealed somatodendritic staining in the CA3 subfield (B,B$_1$), as well as somatic labelling in the dentate gyrus (B$_2$), amygdala (C,C$_1$), caudate putamen (D), thalamus (E), substantia nigra regions (F,F$_1$), hypothalamus (G,H), cerebellum (I,I$_1$) and spinal cord (J,J$_1$). *Abbreviations*: 3V, third ventricle; ARC, arcuate nucleus; amg, amygdala; aq, aqueduct; CA1-3, cornu ammonis 1-3; cc, corpus callosum; CPu, caudate putamen; Ctx, cortex; DG, dentate gyrus; DH, dorsal horn; gl, granular layer; h, hilus; Hp, hippocampus; Hy, hypothalamus; igl, internal granular layer; int, interal capsule; LV, lateral ventricle; Mb, midbrain; ml, molecular layer; pcl, Purkinje cell layer; PVN, paracentricular nucleus; sgz, subgranular zone; sl, stratum lacunosum moleculare; SNc, substantia nigra pars compacta; SNr, substantia nigra pars reticulata; sp, stratum pyramidale; sr, stratum radiatum; Th, thalamus; VH, ventral horn; VTA, ventral tegmental area; wm, white matter. *Scale bars* = 500 µm (C,F); 100 µm (B,D,E,F$_1$,C$_1$,I); 50 µm (B$_1$,B$_2$,G,H,I$_1$,J); 10 µm (J$_1$).

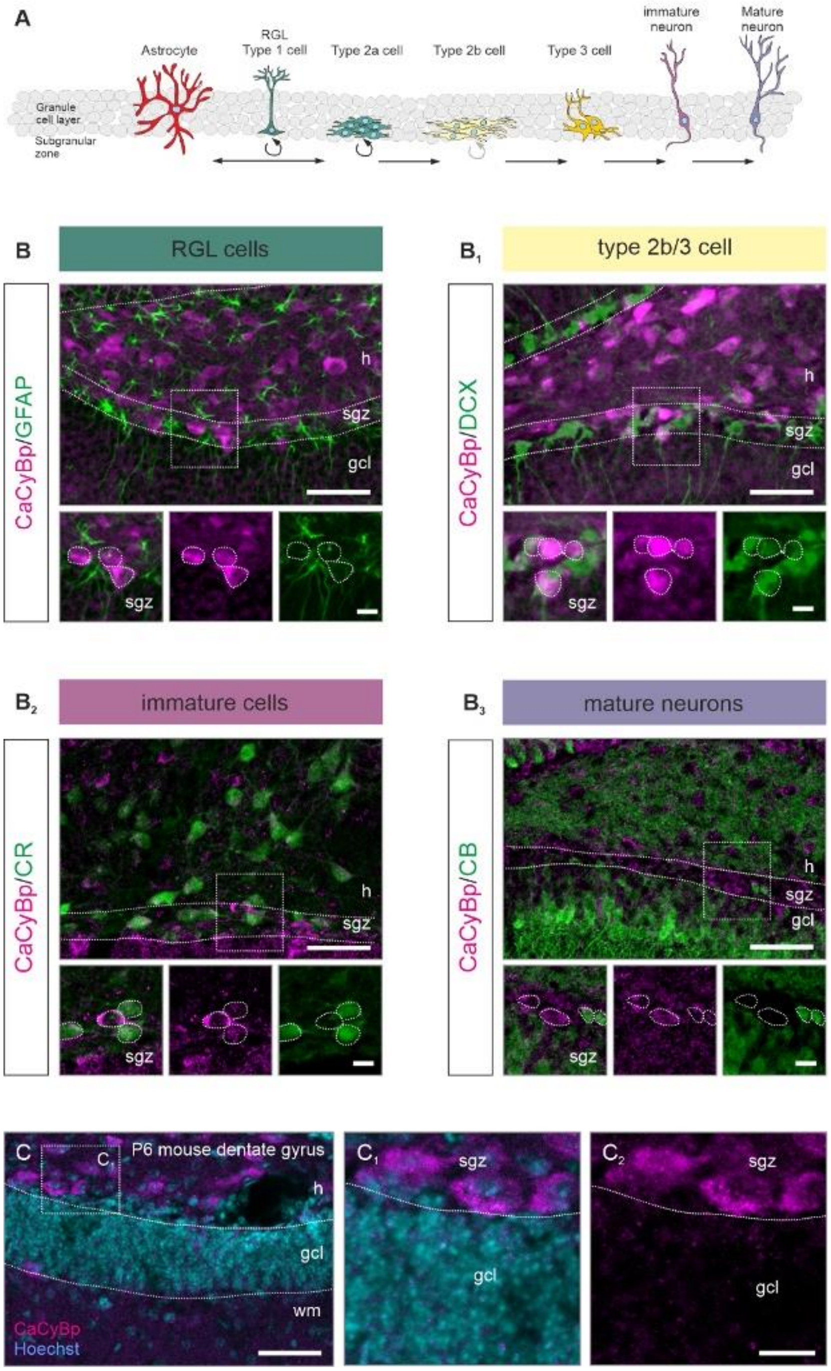


**Supplementary Figure 6. CaCyBp distribution in the mouse dentate gyrus.** (**A**) Schematic of neurogenesis in the adult dentate gyrus (adapted from *Ref*[2]). (**B-B$_3$**) In the subgranular zone, CaCyBp labels new-born doublecortin (DCX)$^+$ neurons (B$_1$) and immature calretinin (CR)$^+$ neurons (B$_2$), but not calbindin$^+$ cells or radial glia (GFAP; B,B$_3$). (**C-C$_2$**) Fluorescent immunohistochemistry for CaCyBp reveals a lack of signal in the granular layer already at postnatal day 6, as compared to E18.5. *Abbreviations*: gl, granular layer; h, hilus; sgz, subgranular zone; wm, white matter. *Scale bars* = 50 µm (B,B$_1$,B$_2$,B$_3$,C), 10 µm (white boxes).

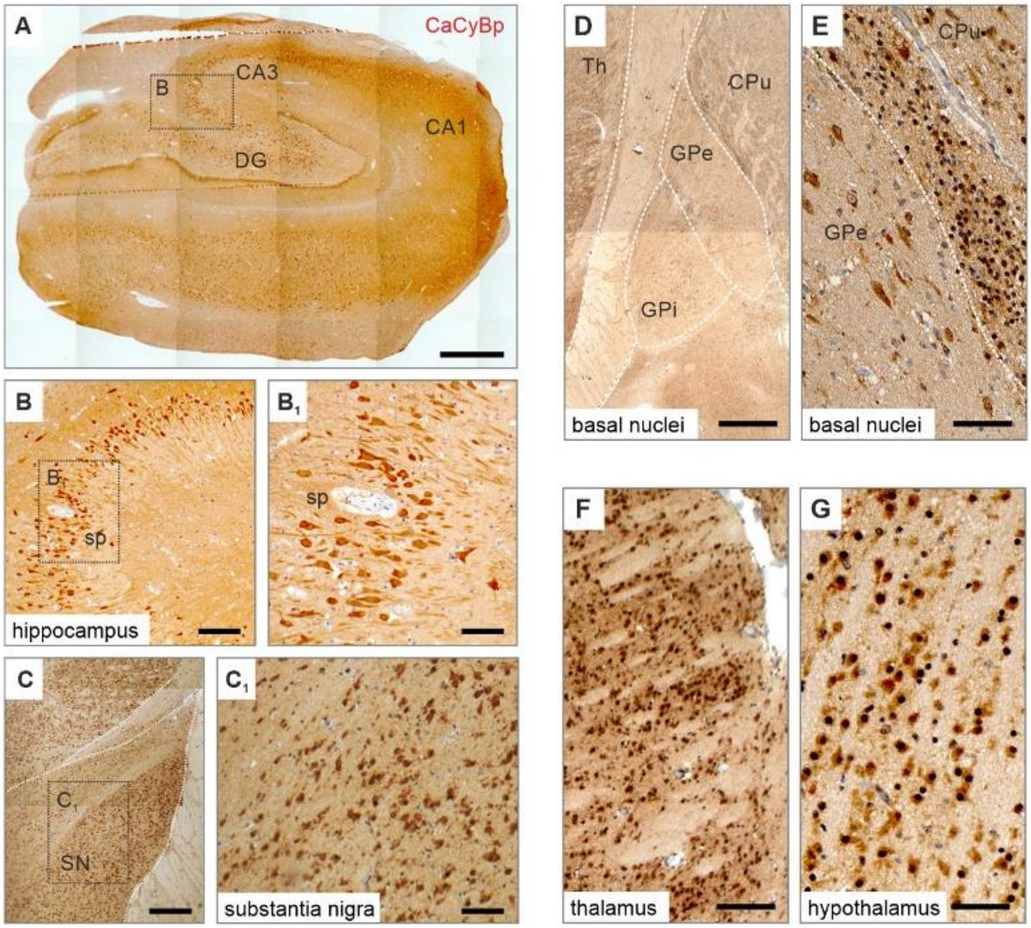


**Supplementary Figure 7. CaCyBp labelling in the adult human brain.** (**A-G**) Immunohistochemistry for CaCyBp in the human adult hippocampal formation Somatodendritic labelling was detected in the pyramidal layer (B,$B_1$), alike in the substantia nigra (C,$C_1$), basal nuclei (D,E), thalamus (F) and hypothalamus (G). *Abbreviations*: CA1-3, cornu ammonis 1-3; CPu, caudate putamen; DG, dentate gyrus; GPe, globus pallidus external; GPi, globus pallidus internal; SN, substantia nigra; Th, thalamus. *Scale bars* = 500 µm (A), 200 µm (C,D,E), 100 µm (B,$C_1$,F,G), 50 µm ($B_1$).

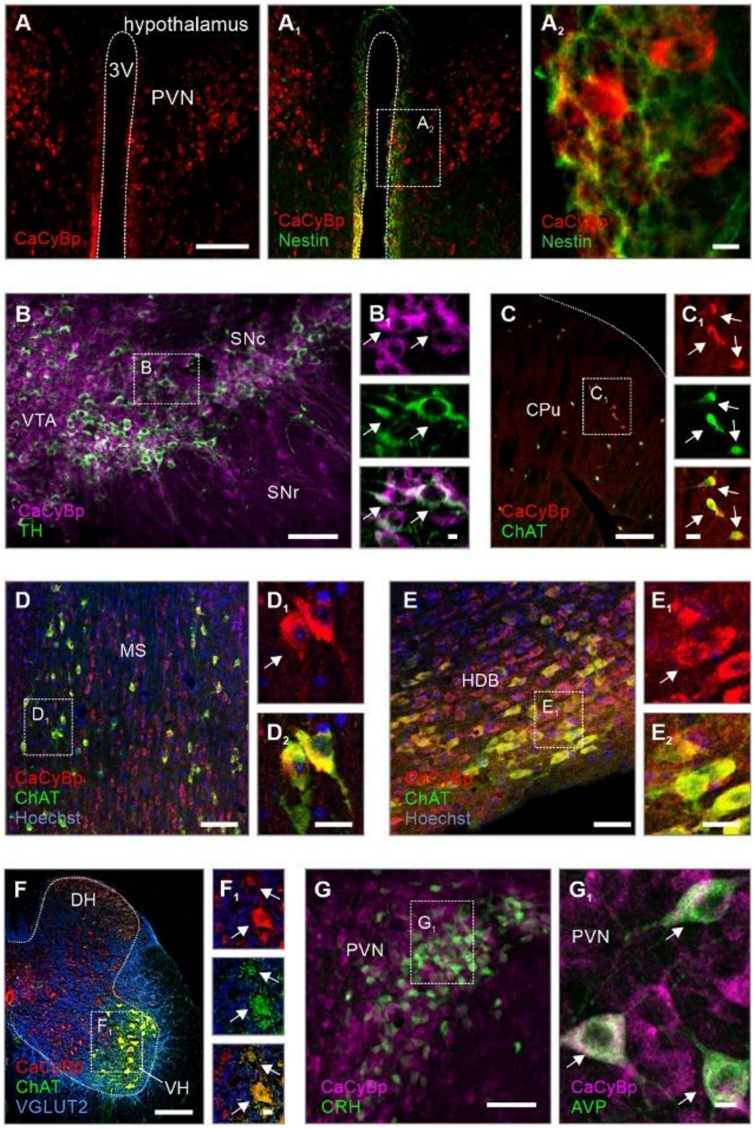


**Supplementary Figure 8. Cell type specificity of CaCyBp expression.** (**A-$A_2$**) Nestin, an ependymal cell marker[3,4], did not colocalize with CaCyBp (hypothalamus is shown). (**B,$B_1$**) A subpopulation of $CaCyBp^+$ cells co-labelled for tyrosine-hydroxylase (TH) in the ventral tegmentum and substantia nigra*pars compacta*. (**C-$F_1$**) CaCyBp also co-localized with choline-acetyltransferase (ChAT) in the dorsal striatum (C,$C_1$), medial septum (D-$D_2$), horizontal diagonal bands (E-$E_2$), and ventral horn of the spinal cord (F,$F_1$). (**G,$G_1$**) In the hypothalamus, CaCyBp was expressed in CRH (G) and AVP-containing cells ($G_1$) of the para-ventricular nucleus. Arrows denote co-localization. *Abbreviations*: 3V, third ventricle; CPu, caudate putamen; DH, dorsal horn; HDB; horizontal diagonal band of Broca; MS, medial septum; PVN, paraventricular nucleus; SNc, substantia nigra pars compacta; SNr, substantia nigra pars reticulata; VN, ventral horn; VTA, ventral tegmental area. *Scale bars* = 100 µm (D,F), 50 µm (A,B,C,E,G), 10 µm ($A_2$,$B_1$,$C_2$,E2,$F_1$,$G_1$).
